# CIRKO: A chemical-induced reversible gene knockout system for studying gene function *in situ*

**DOI:** 10.1101/2022.08.31.506064

**Authors:** Hui Shi, Qin Jin, Fangbing Chen, Zhen Ouyang, Shixue Gou, Xiaoyi Liu, Lei Li, Shuangshuang Mu, Chengdan Lai, Quanjun Zhang, Yinghua Ye, Kepin Wang, Liangxue Lai

## Abstract

Conditional loss and restoration of function are becoming important approaches for investigating gene function. Given that reversible conditional gene knockouts in cells required complicated manipulation, conditional inactivation and reactivation of a gene in primary somatic cells with limited proliferative capacity and in animal models remain difficult to achieve. Here, we first developed a reportable and reversible conditional intronic cassette (ReCOIN), wherein inactivation and reactivation of the gene are mediated via sequential expression of Cre and Flp recombinases, respectively. The expression pattern of the target gene can be monitored by direct visualization. To simply and tightly control temporal expression of the recombinases, on the basis of ReCOIN, we further presented a dual chemical-induced reversible gene knockout system (CIRKO) by insertion of reverse tetracycline transcriptional activator (rtTA) and tetracycline response element (TRE)-controlled Cre and FlpoERT2 recombinases cassettes into *Rosa26* and *Hipp11* loci of cells, respectively, in which transcription termination of the target gene can be induced at a specific stage in the presence of doxycycline, while gene restoration is achieved in the presence of doxycycline and tamoxifen simultaneously. This system provides a simple, rapid, and flexible gene switch for studying gene function *in situ* both *in vitro* and *in vivo*.

**Impact statement:** A novel chemical-induced reversible gene knockout system provides a simple, rapid, and flexible gene switch to facilitate the study of gene function in primary somatic cells *in vitro*, embryos *in vitro* or *in vivo*, and animals *in vivo*.

## Introduction

Conditional loss-of-function approaches have been developed to explore the function of genes essential for embryonic development or investigate the role of specific gene in the specific tissue or specific developmental stage (Housden et al., 2017). The most common strategy used to achieve conditional gene knockout is based on the site-specific recombinase system, such as the Cre/loxP and Flp/FRT systems (Lewandoski, 2001 Yarmolinsky and Hoess, 2015). These recombinases facilitate deletion or inversion of the gene of interest that is flanked by similarly or inversely oriented site-specific recombinase recognition sites, respectively. Manipulating the expression of the recombinase under the control of tissue/cell-specific promoters and at specific time points enables spatiotemporally controlled gene inactivation (Lewandoski, 2001). Temporal control of the recombinase function can be achieved by the Tet-On or estrogen receptor (ER) system only in the presence of two inducers, doxycycline (Dox) and tamoxifen (Tam), respectively (Feil et al., 1997 Schwenk et al., 1998 Utomo et al., 1999). However, these conditional gene inactivation strategies are irreversible. After the loss of gene function, exogenous gene overexpression system is commonly used to rescue the lost gene function (Imperatore et al., 2017). Given the uncertainty of copy number and quantity of expression, the gene overexpression system cannot fully mimic the level of expression of the gene prior to loss of function. To solve this issue, a genetic tool with a capacity to inactivate and reactivate a specific gene *in situ*, which is able to authentically mimic the endogenous expression pattern before inactivation and after restoration is a necessity.

Several previous studies have attempted to conditionally and reversibly regulate the gene expression via site-specific recombinases with different genetic methods, but each of them has limitations. The COIN allele, as an artificial intron, can be used to achieve conditional knockout of target genes, but it is not reversible (Economides et al., 2013). HIT-trap cannot sequentially achieve conditional gene inactivation and restoration (Lu et al., 2021). The orientation of inserted HIT-trap donors can be either forward or inverse. The forward inserted HIT-trap directly inactivates the target gene without the presence of recombinase, although the expression of the knocked-out target gene can be restored by Cre recombinase. Gene knockout with an inverted insertion can be conditionally controlled by Cre recombinase, but it is irreversible. XTR system containing a synthetic fluorescent reporter trap element can achieve gene inactivation by Cre-mediated inversion of an integrated gene-trap reporter, and gene rescue by Flp-dependent deletion of the inverted gene trap, but this system is not suitable for inactivation and restoration of genes with only one exon or any exogenous transgene containing only coding sequence (CDS) (Robles-Oteiza et al., 2015). CRISPR-FLIP, similar to COIN, in which an invertible intronic cassette is designed and inserted into the exon of a specific gene, provides a universal method for bi-allelic conditional and reversible gene knockouts in cells (Andersson-Rolf et al., 2017). However, in CRISPR-FLIP, the DsRed (reporter gene) is placed downstream of the PGK promoter, resulting in expression of DsRed in cells both before and after target gene inactivation. Thus it could not be used to display whether the target gene is inactivated in real-time. Given that conditional and reversible gene knockouts in cells require three rounds of cell transfection and screening, FLIP has merely been applied to the cells with long-term proliferative capacity, such as immortalized cells and pluripotent stem cells, but is not suitable for installing the conditional and reversible knockout allele into primary somatic cells due to their limited expanding ability. Unlike mice and rats, many large animals do not have ES cells with germline competent chimeric capacity, resulting in the inability to construct genetically modified animal models by means of chimeras. The construction of gene-modified large animal models rather relies on somatic cell nuclear transfer (SCNT) approach by using gene-modified primary cells as donor nuclei. Therefore, it is difficult to generate conditional and reversible gene knockout animal models through FLIP approach.

To address the issues above, in this study we designed a reportable and reversible conditional intronic cassette, termed ReCOIN, which consisted of a 5′ splice donor site (5′ss), two oppositely oriented 3′ splice acceptor sites (3′ss), an inverted fluorescent reporter trap element, a puromycin selection cassette, pairs of oppositely oriented *lox*P and *lox*P2272 sites, and a pair of FRT sites. Given the presence of flanking splice sites, ReCOIN can play a role of an artificial intron when inserted into an exon of any target gene, thus not affecting expression pattern of the target gene. In the presence of Cre recombinase, the inverted EGFP reporter and transcription termination signal were reversed, resulting in transcription termination of the target gene and expression of EGFP, which played a role of marker to display inactivation of targeted genes. The inactivation of the targeted genes could be restored via Flp recombinase-dependent deletion of transcription termination signal and could be indicated by disappearance of green fluorescence. Therefore, our ReCOIN system could be used to create primary somatic cells with conditional and reversible knockout elements in endogenous genes. The genetically modified primary somatic cells can be used to clone animal models containing ReCOIN element, thereby providing a new perspective to investigate gene function *in vivo* through inactivation and reactivation of any genes.

The conditional gene inactivation and reactivation allele described above rely on introducing recombinases to cells or animals, which is a complicated and inefficient procedure, especially difficult for primary cells with limited proliferative capacity and large animals such as pigs, dogs and monkeys with long-term reproduction cycles. Dox and Tam-mediated chemical induction of recombinase had been widely used to regulate gene expression temporally (Lewandoski, 2001). Therefore, on the basis of ReCOIN, we developed a dual chemical-induced reversible knockout system (CIRKO) by combining the Tet-On and ERT2 systems with the universal ReCOIN allele. The inactivation and reactivation of gene of interest could be achieved by simple administration with small molecule compounds. Using this system, chemical-induced inactivation and reactivation were achieved in porcine *TP53, LMNA* and *POU5F1* genes. The ReCOIN-based CIRKO system would provide an efficient, rapid, and flexible on-off-on switch of gene to study gene function.

## Results

### Design of the ReCOIN allele

We first designed a conditional conversion gene knockout allele named ReCOIN, which could be universally used for inactivation and reactivation of any gene *in situ* by dual site-specific DNA recombinases. The ReCOIN allele consisted of a *lox*P-flanked inverted gene trap, a puromycin selection cassette, a pair of FRT recognition sites of Flpo recombinase (mammalian codon-optimized Flp), and conserved 5′ss and 3′ss (**Figure 1A; Figure 1-figure supplement 1, 2**). The gene trap comprised 3′ss, coding sequence for EGFP, and ß-globin polyadenylation transcription termination signal (rßglpA) (**Figure 1A; Figure 1-figure supplement 1**). Given that the ReCOIN allele was flanked by 5′ss and 3′ss, it could serve as an exogenous artificial intron after integrating into the exon of target gene. In the process of mRNA splicing, the ReCOIN allele could be spliced out, and the separated left (Exon-L) and right (Exon-R) parts of the exon could be fused together to form a complete exon again, which did not affect the expression pattern of target gene. In the presence of Cre recombinase, the intermediate state with induced inversion of the intervening DNA at either the inverted *lox*P or *lox*P2272 sites were first generated, followed by irreversibly removing the puromycin selection cassette between either two identically oriented *lox*P or *lox*P2272 sites (**Figure 1-figure supplement 1**). During mRNA splicing, the sequences between 5′ss and proximal 3′ss were spliced out, resulting in the fusion of Exon-L and EGFP-polyA, which would inactivate the target gene (**Figure 1A; Figure 1-figure supplement 1**). The EGFP reporter could visualize the cells with inactivation of target gene. Furthermore, Flp recombinase-mediated deletion of the gene trap blocked the EGFP expression and formed the 5′ss-FRT-3′ss artificial intron, which enabled the inactivated target gene to be reactivated (**Figure 1A; Figure 1-figure supplement 1**). In theory, the ReCOIN allele operated as a gene switch that allowed turning any target gene off and on flexibly and monitoring the target gene expression profiling by EGFP-based visualization.

**Figure 1.**
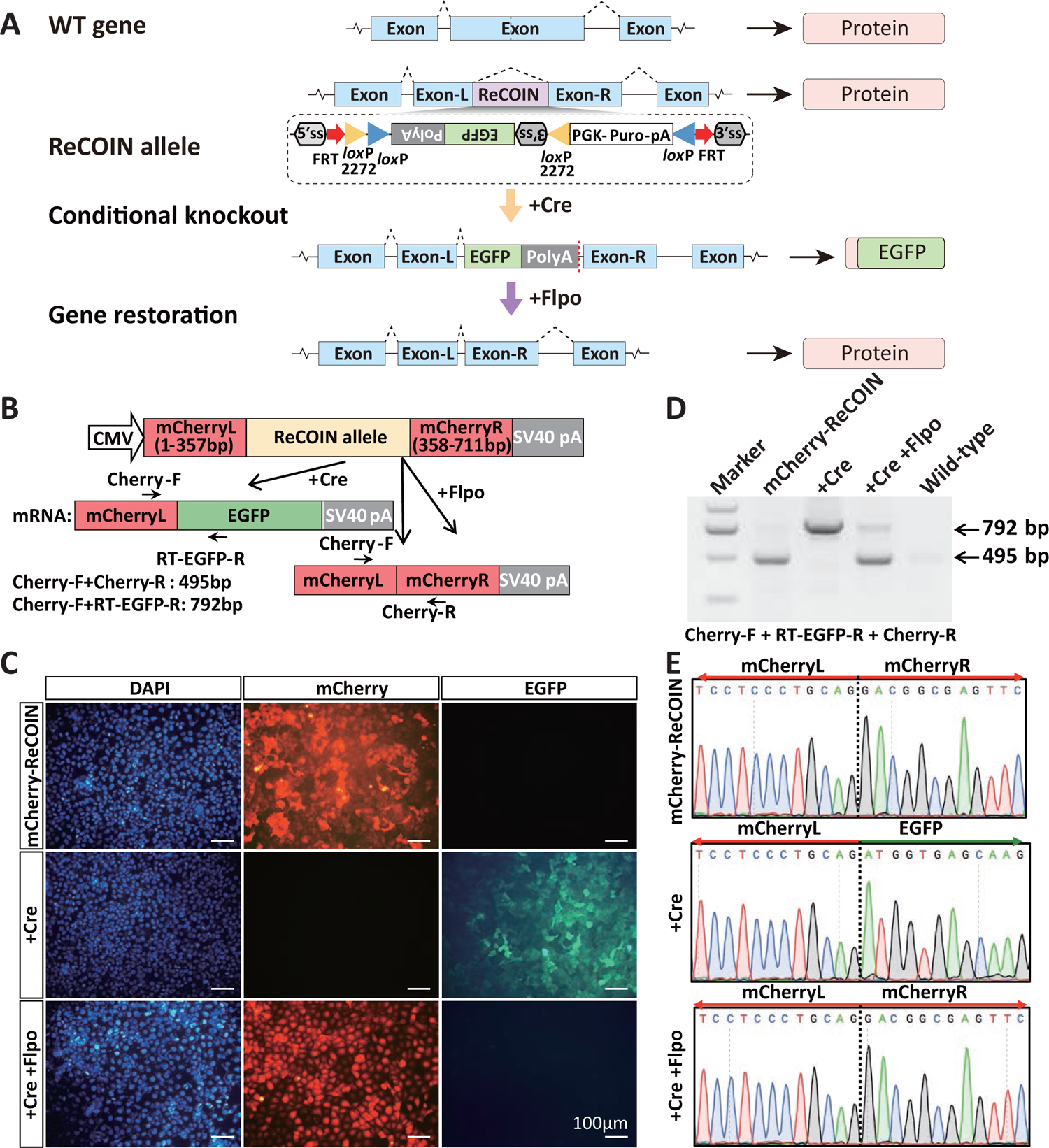
ReCOIN allele for conditional gene inactivation and reactivation. (A) Schematic drawing of the ReCOIN cassette strategy for conditional gene modification. Cre inactivates gene function and Flp restores gene function. Detailed schematic is shown in Figure 1-figure supplement 1. The ReCOIN module is framed by a black dotted box. (B) Schematic of insertion of the ReCOIN cassette in the mCherry gene. (C) Fluorescent images of the sorted mCherry-ReCOIN cells, mCherry-ReCOIN^+Cre^ cells, and mCherry-ReCOIN^+Cre+Flpo^ cells. mCherry protein is expressed in mCherry-ReCOIN cells (top row). After Cre recombination, the mCherry expression is disrupted, and EGFP is expressed (middle row). After sequential transfection of Flpo, only mCherry is expressed (bottom row). (D) RT-PCR analysis of mCherry in mCherry-ReCOIN cells, mCherry-ReCOIN^+Cre^ cells, and mCherry-ReCOIN^+Cre+Flpo^ cells. All detected bands were of the expected size. (E) Sanger sequencing of RT-PCR products in (D).

### Validation of ReCOIN allele at mCherry reporter gene

To validate this system, we initially inserted the ReCOIN allele into the mCherry gene (mCherry-ReCOIN) and evaluated in human embryonic kidney 293 (HEK293) cells. The vector containing the cytomegalovirus (CMV) promoter-driven mCherry-ReCOIN expression cassette (referred to as pCMV-mCherry-ReCOIN) was constructed (**Figure 1B**). Cells were transfected with pCMV-mCherry-ReCOIN vector, pCMV-mCherry-ReCOIN vector+Cre-expressing vector, or pCMV-mCherry-ReCOIN vector+Flpo-expressing vector (referred to as mCherry-ReCOIN, mCherry-ReCOIN+Cre, and mCherry-ReCOIN+Flpo in figures, respectively). Three days post-transfection, under inverted fluorescence microscope, the red fluorescence was observed in cells of the mCherry-ReCOIN and mCherry-ReCOIN+Flpo groups, while the green fluorescence was observed in cells of the mCherry-ReCOIN+Cre group (**Figure 1-figure supplement 1A**). To verify whether precise recombination occurred in these cells by Cre or Flpo recombinase, the cells were subjected to whole mRNA extraction and RT-PCR analysis. As shown in **Figure 1-figure supplement 1**, the ReCOIN allele was correctly spliced at the mCherryL/R junction before inversion and after restoration by Flpo. After Cre-mediated inversion, the expected products of 792 bp spanning the mCherry-L and EGFP were detected (**Figure 1-figure supplement 1B; Figure 1-figure supplement 1-source data**). Sanger sequencing results of these RT-PCR products further revealed the precise junction of mCherry-L and mCherry-R in the mCherry-ReCOIN and mCherry-ReCOIN+Flpo groups (**Figure 1-figure supplement 1C**). Similarly, after Cre-mediated inversion and excision, mCherry-L-EGFP fusion mRNAs were produced by precise fusion of mCherry-L and EGFP, which led to the inactivation of mCherry (**Figure 1-figure supplement 1C**). The mCherry-L and EGFP mRNAs were in-frame and could translate into mCherry-L-EGFP fusion protein, which could visualize the inactivation of mCherry gene.

To further support the idea that precise recombination occurred by Cre and Flpo recombineses, sequential transfection of HEK293 cells were used to substantiate this point. Initially, we integrated pCMV-mCherry-ReCOIN into HEK293 cells by piggyBac transposase and obtained the positive cells expressing mCherry-ReCOIN after fluorescence-activated cell-sorted (FACS) (referred as mCherry-ReCOIN cells in figures) (**Figure 1C; Figure 1-figure supplement 1A, B; Figure 1-figure supplement 1**). Then, mCherry-ReCOIN cells were transiently transfected with Cre recombinase. EGFP expression was observed and EGFP-positive cells were sorted (referred as mCherry-ReCOIN^+Cre^ cells in figures) (**Figure 1C; Figure 1-figure supplement 1A, B; Figure 1-figure supplement 1**). Next, the sorted EGFP^+^ cells were transfected with Flpo recombinase. mCherry was re-expressed and mCherry^+^ cells were sorted (referred as mCherry-ReCOIN^+Cre+Flpo^ cells) (**Figure 1C; Figure 1-figure supplement 1A, B; Figure 1-figure supplement 1**). The mRNA transcripts of sorted cells were collected and performed RT-PCR. RT-PCR and Sanger sequencing analyses showed ReCOIN allele was correctly spliced at the mCherry-L/R junction in mCherry-ReCOIN and mCherry-ReCOIN^+Cre+Flpo^ cells (**Figure 1D, E; Figure 1-source data**). The Cre-mediated inversion showed the expected products of 792 bp and fusion of mCherry-L and EGFP (**Figure 1D, E; Figure 1-source data**). After Flpo-mediated restoration, the expected products of 495 bp spanning the mCherry-L and mCherry-R were detected (**Figure 1D, E; Figure 1-source data**). In addition, to evaluate whether the insertion of ReCOIN allele affects the expression level of mCherry, we transiently transfected HEK293 cells with pCMV-mCherry or pCMV-mCherry-ReCOIN. Flow cytometry assay was performed to detect the proportion of cells expressing mCherry. An average of 48.7% and 45.8% cells expressed mCherry in the pCMV-mCherry and pCMV-mCherry-ReCOIN groups, respectively, and there was no significant difference between the two groups (**Figure 1-figure supplement 1A, B**). Therefore, the insertion of ReCOIN allele barely affects the expression level of mCherry.

The results above demonstrated that the ReCOIN allele had the capacity to flexibly respond to the induction of Cre and Flpo recombinases and that the consequence of induction could be monitored through EGFP-based visualization.

### Validation of ReCOIN allele at a functionally important gene through regulating Cas9 expression in on-off-on manner

The CRISPR/Cas9 system provides a powerful tool for therapeutic genome editing in eukaryotes (Chen et al., 2021, Cox et al., 2015). However, persistent expression of Cas9 was considered to result in genomic instability, off-target effects, and inflammatory responses, which raise serious safety concerns in clinical applications (Chew, 2018, Leibowitz et al., 2021). Therefore, the cleavage activity of Cas9 should be silenced as early as possible once the gene editing was accomplished. To address this issue, we attempted to develop a switchable Cas9 system, named as Cas9-ReCOIN, based on the ReCOIN allele. In this system, the ReCOIN allele was inserted into Cas9 gene, resulting in two fragments, Cas9-L (665 amino acids [aa], 1-665 aa) and Cas9-R (715 aa, 666-1380 aa) (**Figure 2A**). Theoretically, the Cas9-ReCOIN system had cleavage activity similar to wild-type Cas9 because the inserted ReCOIN allele played a role of an intron. The Cas9-ReCOIN nuclease was inactivated in the presence of Cre recombinase, while the cleavage activity of Cas9-ReCOIN was restored in the presence of Flpo recombinase (**Figure 2A**).

**Figure 2.**
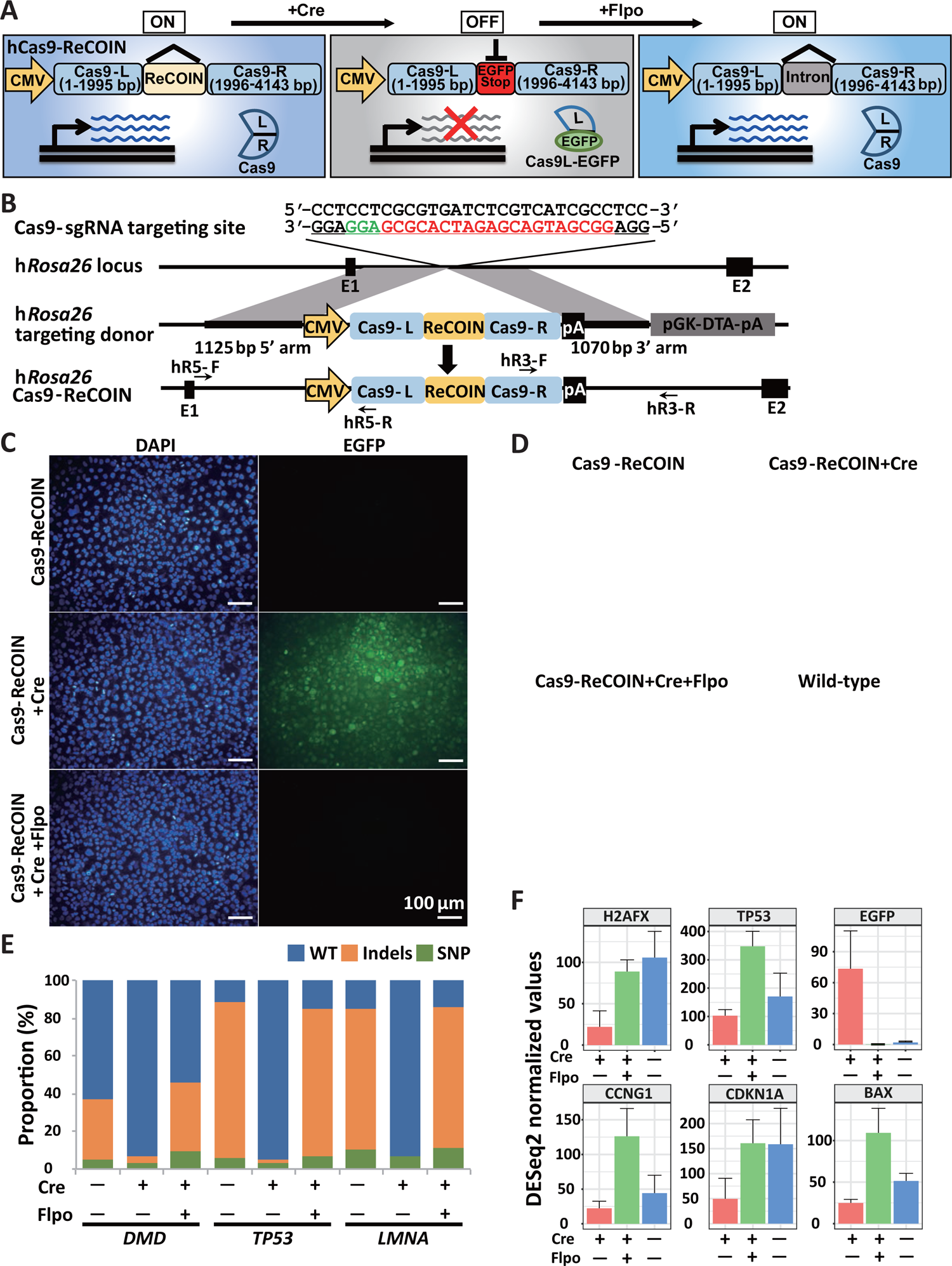
ReCOIN allele facilitates Cre-mediated inactivation and subsequent Flpo-mediated restoration of Cas9. (A) Schematic illustration of Cas9-ReCOIN. The ReCOIN allele functions as an intron and does not disrupt the expression of Cas9. After Cre recombination, the Cas9 expression is disrupted, and EGFP is expressed. Following Flpo recombination, the transcription termination signal is excised and the Cas9 expression is restored. (B) Strategy for introducing Cas-ReCOIN at the hRosa26 locus in HEK293 cells. The sgRNA guide sequence is indicated in red, and the PAM motif is indicated in green. (C) Images of the indicated HEK293 colonies. (D) Detection of EGFP reporter expression in the indicated HEK293 colonies by flow cytometry analysis. (E) Targeting efficiency of DMD, TP53 or LMNA sgRNA in the indicated colonies. (F) Expression levels of H2AFX, TP53, EGFP and genes related to TP53 signaling pathway by transcriptome sequencing.

To verify the hypothesis above, we first tested whether Cas9-ReCOIN could maintain the original enzymatic activity by co-transfecting Cas9-ReCOIN and the EGFP-sgRNA into the HEK293-h*Rosa26*-EGFP cell line, which expressed EGFP under the control of an endogenous h*Rosa26* promoter. Five days post-transfection, we found that the EFGP expression was silent in 50.3% of cells, which was comparable to wild-type Cas9 (59.8%) (**Figure 2-figure supplement 2**). To more conveniently evaluate whether Cas9-ReCOIN could be regulated by recombinases, a HEK293-h*Rosa26*-Cas9-ReCOIN cell line (6#, 1/23, 4.3%), in which the Cas9-ReCOIN allele was integrated into the h*Rosa26* locus of HEK293 cells, was established through puromycin screening and confirmed by 5′ and 3′ junction PCR analysis (**Figure 2B; Figure 2-figure supplement 2A; Figure 2-figure supplement 2-source data 1, 2**). After transfection with Cre recombinase in the HEK293-h*Rosa26*-Cas9-ReCOIN cell line, EGFP-positive colonies (+Cre) were picked up and subcultured to expand population. Flpo recombinase was subsequently transfected into the EGFP-positive cell line, and EGFP-negative colonies were selected (+Flpo). After expansion, genetic characterization confirmed that these selected colonies harbored the modified ReCOIN allele in the *Rosa26* locus as expected (**Figure 2-figure supplement 2B-D; Figure 2-figure supplement 2-source data 3-5**). PCR analysis of these selected colonies showed that Cre and Flpo recombinases could correctly invert and delete the ReCOIN allele within Cas9, respectively (**Figure 2-figure supplement 2E; Figure 2-figure supplement 2-source data 6, 7**). Cre-transfected Cas9-ReCOIN colonies expressed EGFP, while no EGFP expression was observed in Flpo-transfected Cas9-ReCOIN colonies (**Figure 2C, D**). In addition, RT-PCR and Sanger sequencing analyses were performed to confirm expected splicing of the ReCOIN allele during transcription. Results showed that Cas9 and EGFP were joined together only in colonies transfected with Cre recombinase, whereas the two parts of Cas9 that were cleaved by the ReCOIN allele were rejoined together in colonies transfected with Flpo recombinase and in untreated colonies, indicating that the ReCOIN allele was spliced out (**Figure 2-figure supplement 2F**).

We then verified whether the ReCOIN allele could regulate the enzymatic activity of Cas9. The sgRNA targeting three endogenous genomic loci, namely, *DMD*, *TP53*, and *LMNA*, was individually transfected into the cells treated with Cre, Flpo recombinase, and untreated Cas9-ReCOIN cells. The T7 endonuclease I (T7EN1) cleavage assay showed a loss of gene-editing activity of Cas9 in the cell colony transfected with Cre recombinase, whereas cleavage bands were observed in both Flpo-transfected colony and untreated cells with Cas9-ReCOIN allele (**Figure 2-figure supplement 2G-I; Figure 2-figure supplement 2-source data 8-10**). Deep sequencing of the target sites was used to quantify gene-editing activity. Consistent with the T7EN1 cleavage assay, we could hardly observe genome editing in the Cre-transfected colony (**Figure 2E**). Notably, the indel frequencies at the target sites in Flpo-transfected colony were comparable with those in untreated Cas9-ReCOIN cells (36.8% vs. 31.8% for *DMD* gene, 78.0% vs. 82.3% for *TP53* gene, and 74.9% vs. 74.2% for *LMNA* gene) (**Figure 2E**). These results indicated that the insertion of ReCOIN allele into Cas9 allowed flexible regulation of gene-editing activity by Cre and Flpo recombinases. Therefore, Cas9-ReCOIN is a robust inducible genome-editing system. In view of off-target effects and inflammatory responses of constitutive Cas9 systems, it is ideal to control Cas9 expression within a specific time window (Lundin et al., 2020), which has potential to improve the specificity of the endonuclease by reducing activity following on-target modification (Davis et al., 2015Liu et al., 2016). Hence, the ReCOIN allele could be applied to engineer the endonuclease to enable flexible and safe genome editing.

Recent reports showed that Cas9 promotes genomic instability and activates the TP53 pathway (Enache et al., 2020Xu et al., 2020). After we transfected Cre recombinase into the HEK293-h*Rosa26*-Cas9-ReCOIN cell line, the expression of Cas9 was inactivated, which was accompanied with the expression of EGFP (**Figure 2D, F**). The expression levels of *TP53* gene and its downstream target genes, *CCNG1*, *CDKN1A*, and *BAX* on the TP53 signaling pathway, as well as DNA double-strand breaks associated marker gene, γH2AX (alias H2AFX), were down-regulated compared with the untreated h*Rosa26*-Cas9-ReCOIN cell line (**Figure 2F**). When we transfected Flpo recombinase into the Cre-treated cells, Cas9 was reactivated, and the expression levels of *TP53* gene, its downstream target genes, and DNA double-strand breaks associated marker gene were up-regulated again compared with those in the Cre-treated cells (**Figure 2F**). These data demonstrated that the activity of Cas9 nuclease modified by ReCOIN allele could be flexibly regulated, which holds a promise to minimize safety concerns caused by persistent expression of Cas9 in clinical gene therapy.

### Validation of ReCOIN allele at an endogenous *TP53* gene in primary somatic cells and generation of a pig model with ReCOIN allele at *TP53* gene

TP53 is a transcription factor encoded by the *TP53* gene that triggers growth inhibition and apoptotic responses to a range of insults, including DNA damage, stress, and oncogene activation (Martins et al., 2006). Inactivation of TP53 function is a common feature of human tumors and is often associated with increased malignancy, poor survival and resistance to treatment (Vogelstein et al., 2000 Vousden and Lu, 2002). Currently, a variety of *TP53*-modified mouse models have been used to study tumors, including knockout models and induced oncogenic-activating mutation models (Donehower and Lozano, 2009). Genetically engineered mouse models have been widely used for cancer research, but biological differences between humans and mice are obvious. The tumors generated in mice have a higher burden on body weight than those in humans, leading to limitations in utilizing mice to model human cancers and screen anticancer drugs. Given that the anatomy, physiological function, organ development, metabolic regulation, and immune system of pigs are similar to those of humans, pigs are considered ideal large animal models for tumor research (Meurens et al., 2012 Neff, 2019).

TP53 restoration after inactivation has potential for tumor targeted therapy. Therefore, we sought to construct a pig model with reversible gene knockout allele for the study of tumor *in vivo*. To generate the pig model, we targeted the ReCOIN allele to the second exon of p*TP53* gene (**Figure 3A**). The targeting vector containing 988 bp 5′ arm and 947 bp 3′ arm was designed and constructed for homologous recombination. The homologous arms span 1935 bp of the p*TP53* locus containing the exon 2, 3, 4, and the corresponding introns. By inserting the ReCOIN allele into the homologous arms, the transcribed sequences of the second exon of p*TP53* were split into two parts, *TP53*-E2-L (48 bp) and *TP53*-E2-R (26 bp). In addition, one synonymous mutation (GTG > GTA) on the PAM site was introduced in the targeting vector to avoid repetitive digestion. For gene targeting, *TP53*-sgRNA targeting the exon 2 of p*TP53* gene was designed and constructed. The linear targeting vector, *TP53*-sgRNA and Cas9-expressing vector were co-electroporated into porcine fetal fibroblasts (PFFs) derived from 35-day-old Bama mini-pig fetuses. After 10 days of puromycin (400 ng/mL) selection, 127 single cell-derived colonies were selected and expanded to genotype by 5′ and 3′ junction PCR analyses. Genotyping results showed that 22 colonies (22/127, 17.3%) were correctly targeted at the second exon of p*TP53* gene (**Figure 3B; Supplemental Table 1; Figure 3-figure supplement 3; Figure 3-figure supplement 3-source data 1-10**). Further PCR analysis showed that 16 (16/127, 12.6%) were heterozygous knock-in colonies and 6 (6/127, 4.7%) were homozygous knock-in colonies (**Figure 3B; Supplemental Table 1; Figure 3-figure supplement 3; Figure 3-figure supplement 3-source data 1-10**). To validate the functionality of ReCOIN allele in p*TP53* gene, the correctly targeted cell colonies were pooled together and transfected with the Cre recombinase. Three days post-transfection, immunocytochemistry results showed that the EGFP^+^ cells only expressed low levels of TP53 proteins, which might be caused by the incomplete degradation of original TP53 protein within three days (**Figure 3C**). The EGFP^+^ cells were further calculated by flow cytometry. As shown in **Figure 3D**, 36.7% of cells expressed EGFP fluorescence. These EGFP^+^ cells were FACS-sorted and then transfected with Flpo recombinase. Three days after transfection, the Flpo-transfected cells were evaluated by flow cytometry and immunocytochemistry. Compared with the Cre-transfected cells, the Flpo-transfected cells barely expressed EGFP and restored *TP53* expression (**Figure 3C, D**). In addition, whole RNAs were extracted from collected untransfected *TP53*-ReCOIN cells, Cre-transfected cells, and Flpo-transfected cells and used for RT-PCR and Sanger sequencing. The results showed that the expected junction of the left part of *TP53* (*TP53*-E2-L) and EGFP was found in the mRNA transcripts of Cre-transfected cells, whereas normal *TP53* mRNA was transcribed in untransfected *TP53*-ReCOIN cells and Flpo-transfected cells (**Figure 3E, F; Figure 3-source data 1**). These results suggested that the primary porcine somatic cell line with ReCOIN allele integrated into *TP53* was successfully established.

**Figure 3.**
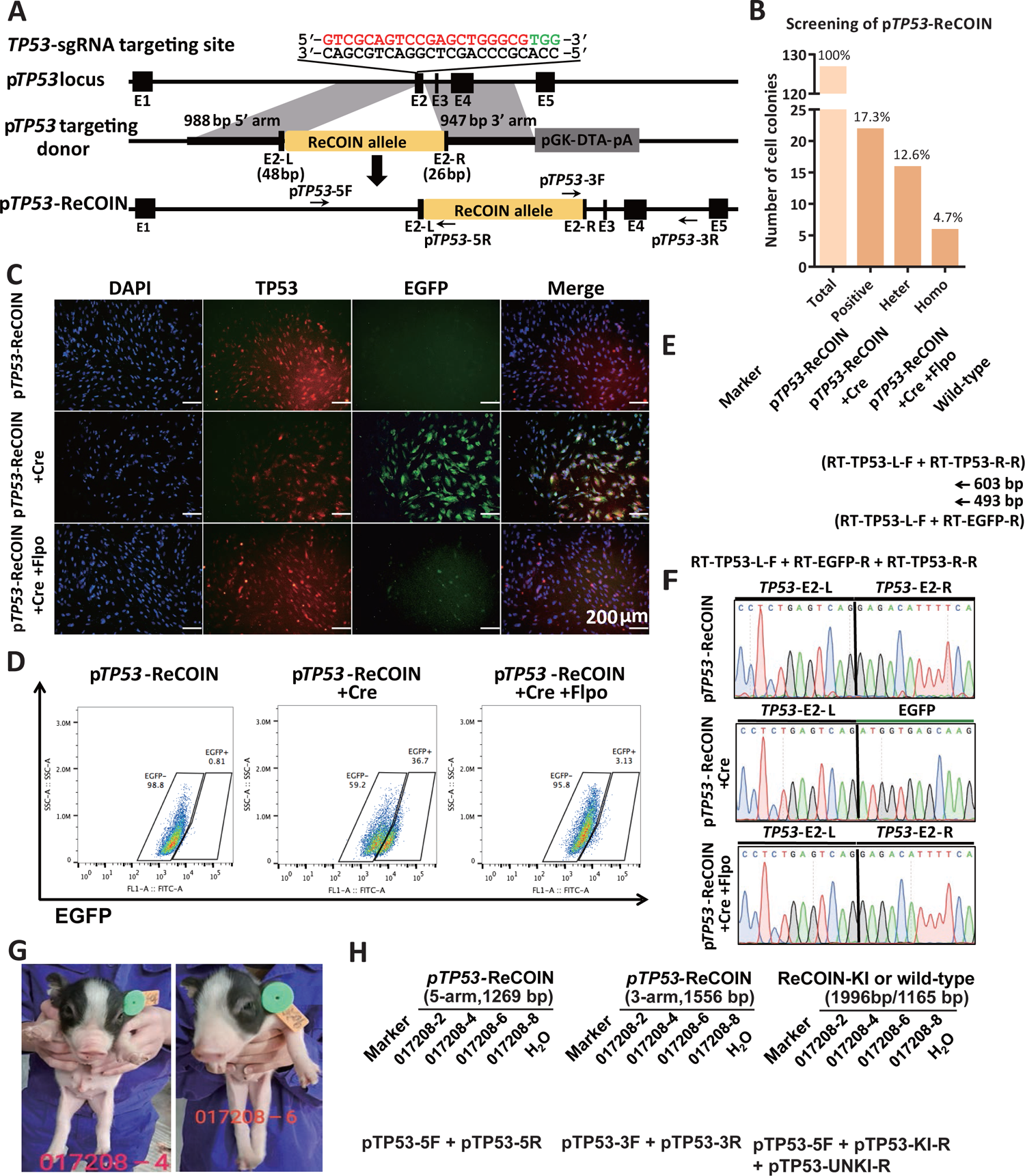
ReCOIN allele facilitate inactivation and subsequent restoration of endogenous p*TP53* gene. (A) Schematic diagram of the strategy for introducing ReCOIN at p*TP53* via homologous recombination in PFFs. The sgRNA is used to target the second exon of p*TP53* gene. The sgRNA guide sequence is indicated in red, and the PAM motif is indicated in green. (B) Histogram of screening the ReCOIN-integrated colonies in endogenous p*TP53* gene. (C) Immunocytochemical staining of TP53 (red) and EGFP (green) in the indicated PFFs. (D) Detection of EGFP reporter expression in the indicated PFFs by flow cytometry analysis. (E) RT-PCR analysis of TP53 in the indicated PFFs. (F) Sanger sequencing of RT-PCR products in (E). (G) Representative photographs of newborn cloned piglets. (H) PCR analysis confirmed correct homologous recombination of the ReCOIN allele at pTP53 locus in ear tissues of the cloned piglets. Three positive piglets had biallelic modifications as detected by PCR. The primers used for genotyping are listed in Supplemental Table 6A.

Four correctly targeted single-cell *TP53*-ReCOIN colonies (C13, C79, C86, and C114) were pooled and used as donor nuclei for SCNT (**Supplemental Table 2**). A total of 785 cloned embryos were reconstructed and transferred into four surrogate pigs (017208, 303306, 305502, and 071706). Of these surrogates, one (017208) was confirmed pregnant by ultrasonography and delivered four cloned piglets (017208-2, 017208-4, 017208-6, and 017208-8) after 114 days of gestation (**Figure 3G; Supplemental Table 2**). PCR genotyping analysis showed that three cloned piglets (017208-2, 017208-6, and 017208-8; 3/4, 75%) carried the ReCOIN allele at the p*TP53* locus (**Figure 3H; Figure 3-source data 2**). Further PCR analysis revealed that all these three piglets were homozygotes and corresponded to the genotype of the C13 and C86 donors (**Figure 3H**). These results demonstrated that live pigs harboring the ReCOIN allele were successfully generated by SCNT, providing a large animal model to study the function of tumor suppressor genes *in vivo*.

### Development of the CIRKO system based on ReCOIN allele

The conditional and reversible gene knockout system described above mediates gene knockout and restoration via Cre and Flpo recombinases, respectively. However, it is difficult to sequentially transfect Cre and Flpo recombinases into primary somatic cells due to the limited cell proliferation ability *in vitro*. Likewise, the sequential delivery of recombinases to animal models via lentiviruses, adeno-associated viruses (AAVs), or lipid nanoparticle is complicated and inefficient. In addition, for large animals, such as pigs, dogs, and monkeys, large-scale virus package and lipid nanoparticle productions are required, which are labor-intensive, time-consuming, and expensive. We took advantage of the chemical-induced system (Tet-On and ERT2 systems) to bypass these obstacles. To temporally regulate gene inactivation and restoration, Tet-On and ERT2 systems were combined with ReCOIN allele to induce recombinases expression (**Figure 4A, B**). The reverse tetracycline transcriptional activator (rtTA) and tetracycline response element (TRE)-controlled Cre and FlpoERT2 recombinases expression cassettes were inserted into p*Rosa26* and p*Hipp11* loci by homologous recombination, respectively (**Figure 4A**). In the absence of exogenous inducers, neither Cre nor Flpo recombinase was expressed, indicating no leaky expression of the recombinase in the system (**Figure 4-figure supplement 4A, B**). In the presence of Dox, Cre and FlpoERT2 were expressed (**Figure 4-figure supplement 4A, B**). Given the inability of FlpoERT2 to enter the nucleus, only Cre recombinase could invert the ReCOIN allele and mediate specific gene inactivation (**Figure 4C**). When Dox and 4-OHT were simultaneously administrated, Flpo could enter the nucleus and induce specific gene restoration (**Figure 4D; Figure 4-figure supplement 4A, B**). The CIRKO system enabled flexible and rapid gene inactivation and restoration, promising a wide range of applications in primary cells *in vitro* and animals *in vivo*.

**Figure 4.**
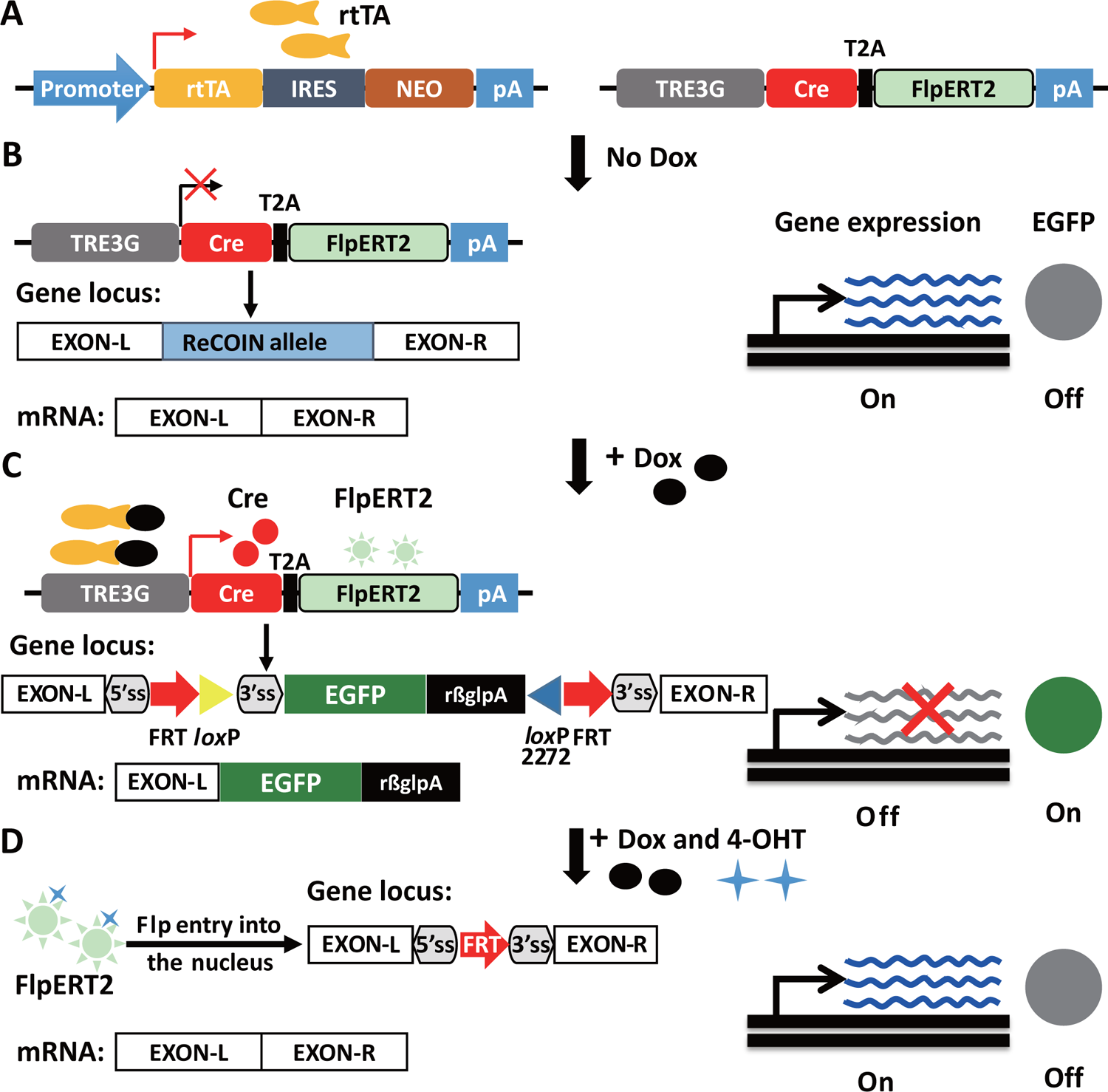
Schematic of CIRKO for chemical-induced reversible gene knockout. (A) Tet-On inducible system was used to drive the expression of Cre recombinase and FlpERT2 fusion. (B) When there is no Dox stimulation, the recombinase is not expressed, and the ReCOIN allele inserted into an exon of the target gene like an artificial intron and does not affect target gene expression. (C) In the presence of Dox, Cre and FlpoERT2 are expressed, at which point FlpoERT2 couldn’t enter the nucleus. Cre inverts the EGFP reporter and the transcription termination signal, resulting in transcription termination of the target gene and visualization of the cells in which the target gene is inactivated by EGFP labeling. (D) In the presence of both exogenous inducers Dox and 4-OHT, Flpo enter the nucleus and induce deletion of the transcription termination signal, thereby leading to gene reactivation.

### Full chemical-induced knockout and restoration of endogenous genes in primary cells and embryos by using the CIRKO system

We next validated whether the CIRKO system enabled inactivation and reactivation of endogenous genes in primary somatic cells by chemical induction. In our group, we have established a PFF cell line with integrated rtTA at the *Rosa26* locus (p*Rosa26*-rtTA)(Jin et al., 2022). The TRE-controlled Cre and FlpoERT2 recombinases expression cassette was inserted into an open intergenic region, the *Hipp11* locus in the PFFs with p*Rosa26*-rtTA (**Figure 5A**). The targeting vector for homologous recombination in the *Hipp11* locus contained 911 bp 5′ homologous arm, 1088 bp 3′ homologous arm, the TRE3G-controlled Cre and FlpoERT2 recombinases expression cassette, and blasticidin resistance gene expression cassette. To achieve chemical-induced p*TP53* gene inactivation and reactivation, the linear p*Hipp11* and p*TP53* targeting vectors, Hipp11-sgRNA, *TP53*-sgRNA, and hCas9-expressing vectors were co-electroporated into p*Rosa26*-rtTA PFFs. After 10 days of selection with puromycin (400 ng/mL) and blasticidin (3 µg/mL), 61 single cell-derived colonies were selected and identified by PCR analysis (**Figure 5B**). The PCR results showed that 9 single cell-derived colonies (CIRKO-p*TP53*-3, 8, 13, 16, 18, 22, 31, 35, and 39) simultaneously contained the ReCOIN allele and TRE3G-controlled two recombinases (Cre and FlpoERT2) expression cassette at p*TP53* and p*Hipp11* loci, respectively (**Figure 5B; Supplemental Table 3; Figure 5-figure supplement 5A, B; Figure 5-figure supplement 5-source data 1-5**). Among these colonies, 6 colonies (CIRKO-p*TP53*-3, 13, 22, 31, 35, and 39) were homozygous knock-in at the exon 2 of p*TP53* gene (**Figure 5B; Supplemental Table 3; Figure 5-figure supplement 5A, B; Figure 5-figure supplement 5-source data 1-5**). The homozygous colonies were treated with Dox for 6 days, followed by simultaneous Dox and 4-hydroxytamoxifen (4-OHT) treatments for additional 3 days (**Figure 5-figure supplement 5A**). The treated cells were collected and analyzed on Day 6 and 9 by PCR and flow cytometry. After 6 days, nearly 50% of the CIRKO-p*TP53* PFFs were EGFP positive as expected (**Figure 5-figure supplement 5B, C**). After Dox and 4-OHT treatments, green fluorescence was gradually disappeared (**Figure 5C, Figure 5-figure supplement 5, 4**). Flow cytometry results showed that about 75.7% CIRKO-*TP53* cells expressed the green fluorescence after 9-days Dox treatment (**Figure 5C**). After simultaneous Dox and 4-OHT treatments for 3 days, the proportion of EGFP^+^ cells was reduced to 10.3% (**Figure 5C**). Immunofluorescence analysis further revealed that the expression of endogenous *TP53* gene was disrupted after Dox treatment while was reactivated after simultaneous Dox and 4-OHT treatments (**Figure 5D**). PCR analysis exhibited that in the CIRKO-*TP53* cells, Dox treatment efficiently and precisely inverted the ReCOIN allele, and subsequent Dox and 4-OHT treatments deleted the transcription termination signal (**Figure 5E; Figure 5-source data 1**). No band with a size of 2747 bp was observed without Dox treatment, indicating no leaky expression of Cre. Likewise, no band with a size of 1342 bp was observed in the absence of 4-OHT treatment, indicating that 4-OHT tightly controlled Flpo incorporation into the nucleus (**Figure 5E; Figure 5-source data 1**). These results suggested that the CIRKO system enabled flexible inactivation and reactivation of a functional gene by sequential chemical induction.

**Figure 5.**
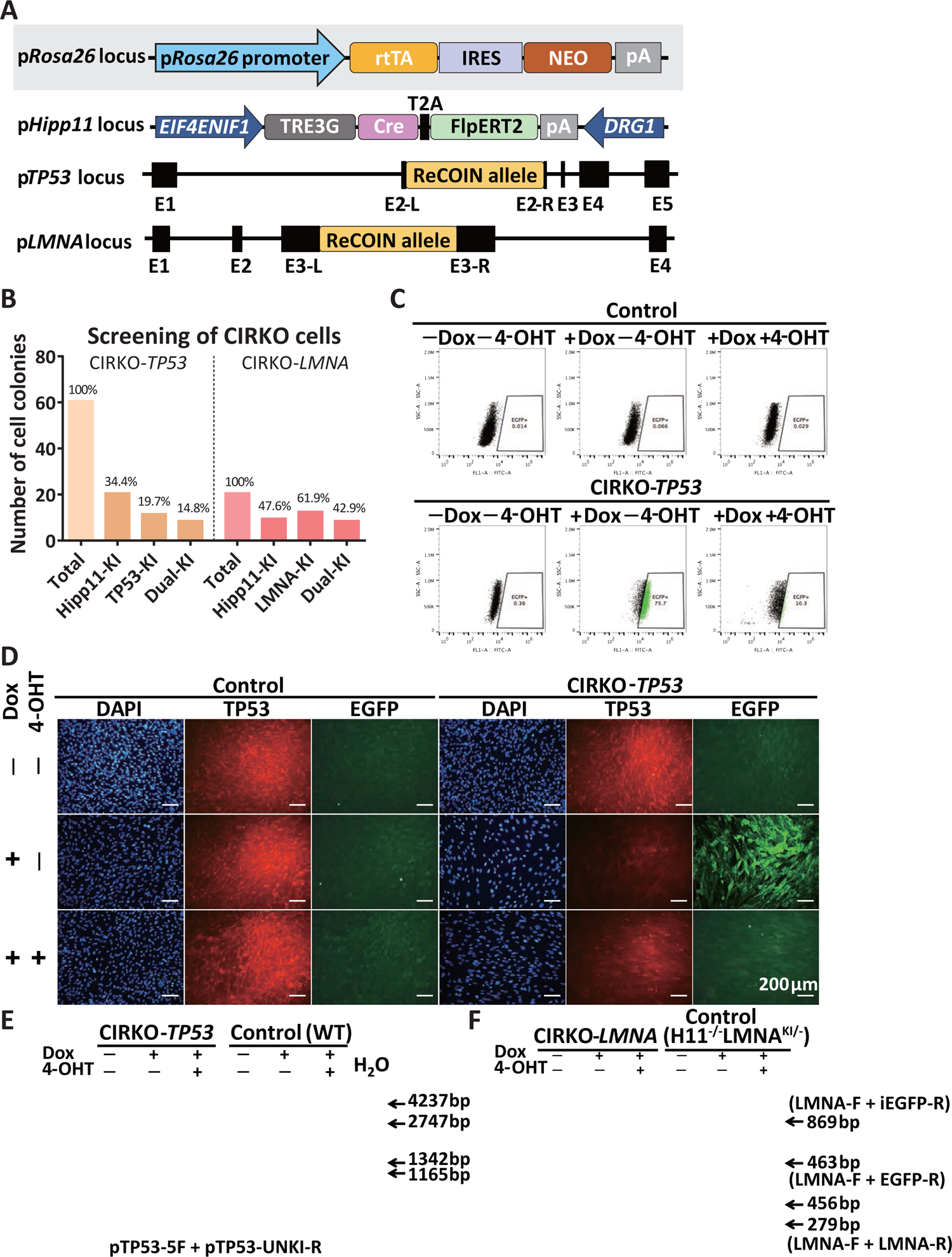
CIRKO facilitates full chemical-induced gene inactivation and subsequent restoration of endogenous genes in primary cells. (A) Schematic diagram of the strategy to generate CIRKO-p*TP53* and CIRKO-p*LMNA* PFFs via homologous recombination. (B) Histogram of screening the CIRKO-integrated colonies in endogenous p*TP53* and p*LMNA* gene. (C) Detection of EGFP reporter expression in the indicated CIRKO-p*TP53* PFFs by flow cytometry analysis at day 9 after treatment. Wild-type PFFs are used as controls. Triplicated experiments for the PFF clone were performed as shown in Figure 5-figure supplement 5. (D) Images of TP53 (red) and EGFP (green) expression in the indicated CIRKO-p*TP53* PFFs as determined by immunocytochemistry. Wild-type PFFs are used as controls. (E) PCR analysis of p*TP53* locus in the indicated CIRKO-p*TP53* PFFs. (F) PCR analysis of p*LMNA* locus in the indicated CIRKO-p*LMNA* PFFs.

To validate the universality of chemical-induced inactivation and reactivation of endogenous genes in primary cells by using the CIRKO system, two other porcine endogenous genes, *LMNA* and *POU5F1*, were chosen as target sites to integrate the ReCOIN allele. Mutations within the *LMNA* gene encoding Lamin A/C would exhibit premature senescence phenotypes at the cellular level and result in aging-related diseases at the individual level (Genschel and Schmidt, 2000). Disruption of *POU5F1 (OCT4)* gene expression would disturb the pluripotency of embryonic stem cells and affect the development of late blastocysts in the mouse (Frum et al., 2013Nichols et al., 1998). However, the developmental phenotype of porcine preimplantation embryos resulting from reversible *POU5F1* perturbation had not been well clarified due to a lack of methods to flexibly perturb *POU5F1* gene expression related to early lineage specifications in porcine embryos.

The ReCOIN allele was inserted into the locus between 5′ and 3′ homologous arms to split the exon 3 of p*LMNA* gene into *LMNA*-E3-L (453 bp) and *LMNA*-E3-R (122 bp), and the exon 1 of p*POU5F1* gene into *POU5F1*-E1-L (237 bp) and *POU5F1*-E1-R (171 bp) (**Figure 5A, 6B; Figure 5-figure supplement 5A; Figure 6-figure supplement 6A**). Similarly, p*Rosa26*-rtTA PFFs were co-transfected with the linear targeting vectors, sgRNAs and Cas9-expressing vectors and plated on the 10 cm culture dishes for clonal isolation. For p*LMNA* gene, 21 single cell-derived colonies were analyzed by PCR, 9 of which (CIRKO-p*LMNA*-2, 4, 5, 8, 10, 12, 13, 15, 16; 9/21, 42.9%) harbored simultaneous insertion of the ReCOIN allele and TRE3G-controlled Cre and FlpoERT2 recombinases expression cassette in the exon 3 of p*LMNA* gene and p*Hipp11* loci, respectively (**Figure 5B; Supplemental Table 3; Figure 5-figure supplement 5B; Figure 5-figure supplement 5-source data 1-3**). Similar to that in CIRKO-p*TP53* cells, untreated, Dox-treated, and Dox + 4-OHT-treated CIRKO-p*LMNA* cells were collected to extract whole DNAs, followed by detection of recombinase-mediated DNA recombination. The PCR results showed that desired DNA bands with length of 463 bp were amplified in the genome of CIRKO-p*LMNA* cells with Dox treatment (**Figure 5F; Figure 5-source data 2, 3**). Likewise, the expected 456 bp bands were found in CIRKO-p*LMNA* cells with simultaneous Dox and 4-OHT treatments, suggesting that precise regulation of p*LMNA* gene disruption and restoration could be achieved by the CIRKO system (**Figure 5F; Figure 5-source data 2, 3**). For p*POU5F1* gene, 9 single cell-derived colonies (CIRKO-p*POU5F1*-3, 12, 13, 15, 19, 28, 32, 35, 36; 9/36, 25.0%) with double-gene knock-in at the p*POU5F1* and p*Hipp11* loci were identified (**Supplemental Table 3; Figure 6-figure supplement 6B, C; Figure 6-figure supplement 6-source data 1-4**). Three of them (CIRKO-p*POU5F1*-3, 19, 32) harbored homozygous ReCOIN allele at the p*POU5F1* locus and they were pooled for further analysis. To well clarify the developmental phenotype of porcine pre-implantation embryos resulting from *POU5F1* perturbation, we performed *POU5F1* perturbation during embryonic development or cellular level. Initially the untreated, Dox-treated, and Dox + 4-OHT-treated CIRKO-p*POU5F1* cells were used as nuclear donors for SCNT (**Figure 6-figure supplement 6A**). No significant differences were observed in blastocyst rate and cell number per blastocyst in the groups of reconstructed embryos from untreated, Dox-treated, and Dox + 4-OHT-treated CIRKO-p*POU5F1* cells (**Figure 6-figure supplement 6B-D; Figure 6-figure supplement 6; Supplemental Table 4**). The PCR products with specific primers were used to further investigate whether chemical-mediated recombination of the ReCOIN allele occurred in these reconstructed embryos. The 738 bp band for embryos from untreated cells, the 324 bp PCR band for embryos from Dox-treated cells, and the 357 bp band for embryos from Dox and 4-OHT-treated cells were observed as expected (**Figure 6-figure supplement 6E; Figure 6-figure supplement 6-source data 1-3**). The expression of POU5F1 and EGFP proteins was evaluated by immunofluorescent staining. The results showed that the expression of POU5F1 was faint by Dox treatment, whereas the expression of EGFP was observed. After Dox and 4-OHT treatments, the restoration of POU5F1 expression was found in the reconstructed CIRKO-p*POU5F1* embryos (**Figure 6-figure supplement 6F**). To take full advantage of the CIRKO system, we performed reversible knockout of p*POU5F1* at the embryonic level. The CIRKO-p*POU5F1* cells were used as nuclear donors for SCNT and the reconstructed embryos were treated with Dox at 4-cell stage while treated with Dox and 4-OHT at 8-cell stage (**Figure 6A**). The reconstructed embryos treated with Dox alone or without any inducers, respectively were used as controls (**Figure 6A**). Consistent with the administration of inducers at the cellular level, followed by SCNT, there were no significant differences in blastocyst rate and cell number per blastocyst among the three groups of reconstructed embryos (**Figure 6C-E; Figure 6-figure supplement 6; Supplemental Table 5**). PCR analyses showed that expected DNA bands with length of 324 bp were observed in the Dox-treated group (**Figure 6-figure supplement 6; Figure 6-figure supplement 6-source data 1-3**). No band with a size of 738 bp was observed with Dox treatment, displaying high inversion efficiency. Likewise, the expected 357 bp bands were found in the reconstructed embryos with simultaneous Dox and 4-OHT treatments, suggesting that restoration of p*POU5F1* gene could be achieved by the CIRKO system (**Figure 6-figure supplement 6; Figure 6-figure supplement 6-source data 1-3**). Notably, there was a band with a size of 1762 bp was observed with simultaneous Dox and 4-OHT treatments in some blastocysts, indicating that incomplete restoration (**Figure 6-figure supplement 6; Figure 6-figure supplement 6-source data 1-3**). Immunofluorescence analysis further showed that the expression of p*POU5F1* was disrupted after Dox treatment while was reactivated after simultaneous Dox and 4-OHT treatments (**Figure 6F**). The results above indicated that, comparing with CRISPR/Cas9 mediated directly editing embryos, CIRKO system enabled real-time and reversible genetic modification in embryo level. Therefore, it provided a universal and rapid strategy for studying the function of genes in pre-implantation embryo level, and potentially in animal level when administrated with different inducers in different time. The established CIRKO system was an effective approach for inactivation and reactivation of endogenous genes in primary somatic cells and embryos. The genetically modified primary somatic cells could be used to clone animal models containing CIRKO elements, which would provide a new perspective to investigate gene function *in vivo* through inactivation and reactivation of any gene.

**Figure 6.**
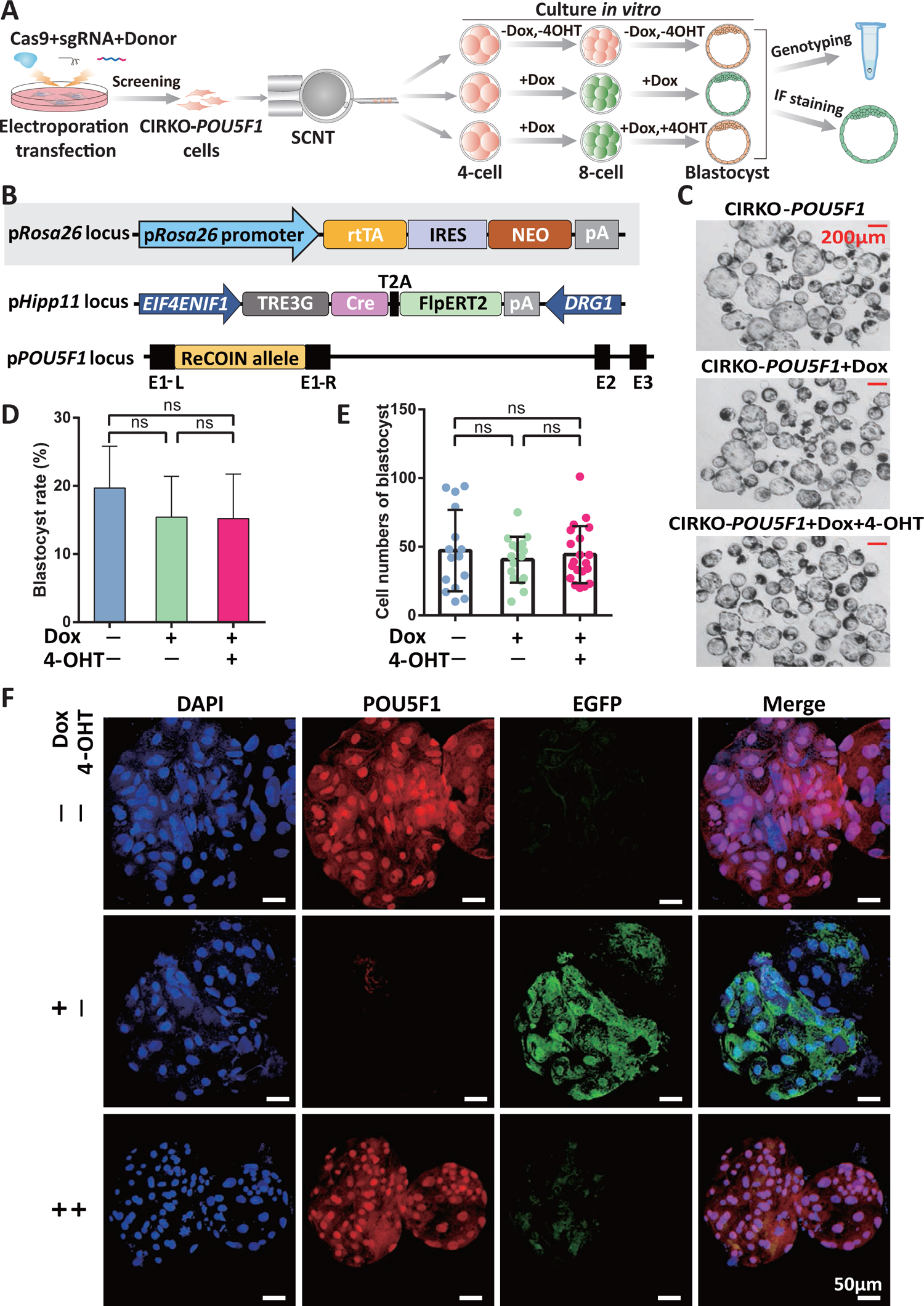
CIRKO facilitates full chemical-induced gene inactivation and subsequent restoration of *POU5F1* in embryos. (A) Experimental procedure to perform reversible knockout of p*POU5F1* at the embryonic level. (B) Schematic diagram of the strategy to generate CIRKO-p*POU5F1* PFFs via homologous recombination. (C) Representative images of untreated (top row), Dox-treated (middle row), and Dox+4-OHT treated (bottom row) CIRKO-p*POU5F1* reconstructed embryos at day 6. (D) Blastocyst rates of untreated, Dox-treated, and Dox+4-OHT treated CIRKO-p*POU5F1* reconstructed embryos. The data are presented as the mean ± SD of four independent experiments. Kruskal-Wallis test was used to calculate *p* values. ns indicates not significant. Source data is available in Supplemental Table 5. (E) Total cell numbers of individual untreated, Dox-treated, and Dox+4-OHT treated CIRKO-p*POU5F1* blastocyst. The data are presented as the mean ± SD. Sample sizes were n=15, 16 and 20 for untreated, Dox-treated, and Dox+4-OHT treated CIRKO-p*POU5F1* reconstructed blastocysts, respectively. Kruskal-Wallis test was used to calculate *p* values. ns indicates not significant. Source data is available in Figure 6-figure supplement 6. (F) Representative confocal images of POU5F1 (red) and EGFP (green) expression in the untreated, Dox-treated, and Dox+4-OHT treated CIRKO-p*POU5F1* blastocysts.

## Discussion

In this study, we first devised a reportable and reversible conditional intronic cassette (ReCOIN) that enabled gene inactivation and reactivation via sequential expression of Cre and Flpo recombinases, respectively. In the ReCOIN allele, we placed the puromycin selection cassette in the downstream of PGK promoter for cell screening. Instead of PGK promoter as that in FLIP (Andersson-Rolf et al., 2017), the promoter of target gene would drive the expression of inverted EGFP and PolyA. EGFP is expressed specifically in cells in which the target gene has been inactivated after Cre recombinase treatment, thereby allowing direct visualization of the cells with inactivated gene and making it convenient to trace and pick up the targeted cells both *in vivo* and *in vitro*, as well as to assess the efficiency of inactivation. Reversible gene knockout can be achieved easily by ReCOIN. The general rules for using ReCOIN are splitting the exon in-frame and splitting the front exon to disrupt more transcription. In addition, the cleavage site obeys the conservative splicing sequence (such as AG/GT, “/” denotes the cleavage site) is preferred.

We verified the validity of the ReCOIN element in the mCherry reporter gene. In addition, we took advantage of this element to conditionally and reversibly regulate the expression of an important functional gene, i.e., Cas9 through “on-off-on” manner. As an exogenous gene in the cells, Cas9 maintains its editing activity after being inserted with the ReCOIN element (on). When Cre recombinase was delivered to cells, Cas9 gene was inactivated (off), and successfully reduced expression levels of *TP53* gene, its downstream target genes on TP53 signaling pathway, and a DNA double-strand break-associated marker gene. When necessary, the inactivated Cas9 could be reactivated by Flpo recombinase and completely restored its cleavage activity (on), which was confirmed by using *DMD*, *TP53*, and *LMNA* as genes of interest. Limiting the editing activity of Cas9 within a specific time window while timely inactivating it outside the time window can reduce off-targets to some extent (Davis et al., 2015Liu et al., 2016). Flexibly regulating the cleavage activity of Cas9 nuclease modified by the ReCOIN allele holds a promise to minimize safety concerns caused by persistent expression of Cas9 protein during CRISPR/Cas9-mediated gene therapy.

Furthermore, the primary somatic cells with ReCOIN allele could be picked by simple visualization of the green fluorescence and used as nuclear donors to make cloned animal models with conditional and reversible knockout element. This system is particularly important for large animals. As far, generation of gene-edited large animal is mainly based on SCNT or direct embryo injection approaches, which are inefficient, expensive, and laborious (Maynard et al., 2021). Therefore, production of conditional and reversible gene knockout large animals remains a challenge due to serial gene targeting and serial rounds of SCNT are needed. As a large animal, pigs are ideal animal models to explore gene function because pigs are closer to humans than rodents in embryonic development, anatomy, physiological systems, and metabolic regulation (Fan and Lai, 2013Zhang et al., 2021). With the ReCOIN system, we successfully generated a pig model, in which endogenous porcine *TP53* could be conditionally and reversibly knocked out. This pig model allows one-step generation of conditional and reversible *TP53* gene knockouts and enables to study tumor suppressor gene function in large animals closer to humans.

To overcome complicated and inefficient procedure for introducing recombinase into cells and animals, by taking the advantage of ReCOIN, we further developed a dual chemical-induced reversible knock out system by combining Tet-On and ERT2 systems with ReCOIN allele. The conditional inactivation and reactivation of gene of interest could be achieved by simple administration with small molecule compounds, allowing flexible regulation of endogenous gene knockout and restoration through drug induction. The CIRKO system can bypass the barrier for delivering recombinases through viruses, which holds promise for widespread application in primary cells *in vitro* and animals *in vivo*. The validity of the CIRKO system was verified on the tumor suppressor gene (p*TP53*), pluripotency-related gene (p*POU5F1*), and aging-related gene (p*LMNA*) in porcine fetal fibroblasts. The primary somatic cells with ReCION, Tet-On and ERT2 elements could be used to generate embryos and animal models for direct investigation of gene functions both *in vitro* and *in vivo*.

The ReCOIN-based CIRKO system established in this study, which has evident advantages, including flexible design, direct visualization of gene expression patterns, removal of drug selection cassette in one step, full chemical-induced reversible gene inactivation, and versatility to study the function of any gene with one or more exons, can be used to study gene function in different biological contexts and is more conducive to studying mechanisms *in vivo* than traditional genetic tools. Given that the CIRKO system allows reversible gene knockout directly via small molecule compounds, it provided a platform to study function of genes related to cancers, pluripotency, development, and metabolism in large levels, such as pigs, dogs, and even in non-human primates with long-term reproduction cycles which have more similarities to humans in evolutionary distance than small animals such as rodents (**Figure 7**). The expected application prospects are as follows: (1) CIRKO can be targeted into tumor suppressor genes to investigate the mechanisms relevant to tumor initiation and progression. (2) CIRKO can be targeted into pluripotency genes to investigate the mechanisms underlying cell fate regulation during reprogramming. (3) CIRKO can be targeted into genes vital for embryonic development to establish live large animal disease models and allow subsequent therapeutic intervention. (4) CIRKO can be exploited to inactivate a putative drug target transiently, allowing prediction of the therapeutic effect and unforeseen complications of the drug.

**Figure 7.**
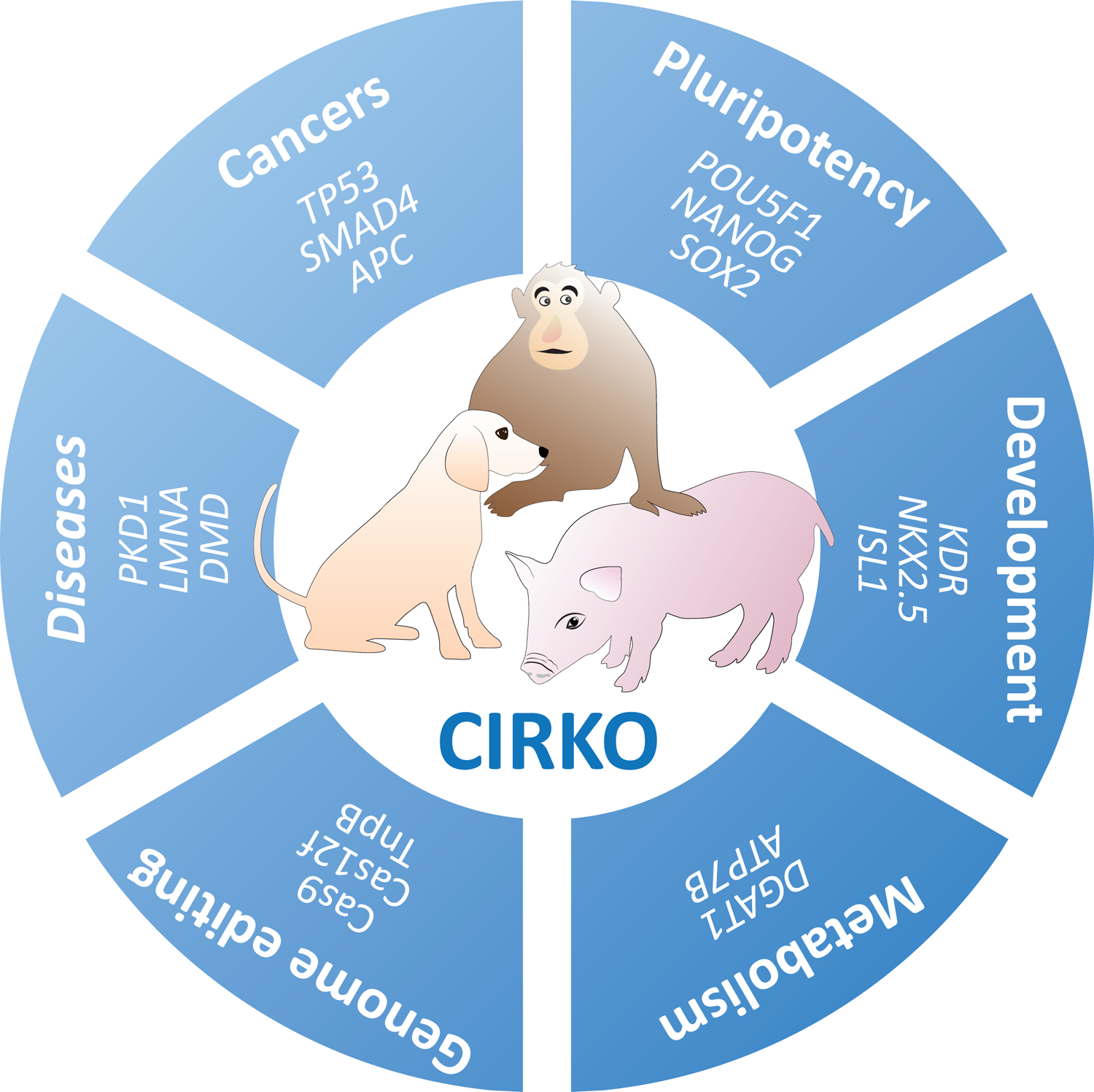
Application prospects of CIRKO system in large animals.

In summary, the conditional and reversible gene knockout system established in this study provides a simple, rapid, and flexible gene switch to facilitate the study of gene function in primary somatic cells *in vitro*, embryos *in vitro* or *in vivo*, and animals *in vivo*.

## Materials and Methods

### Animals

Large white pigs and Bama miniature pigs under conventional housing conditions, maintained at the large animal facility of Guangzhou Institutes of Biomedicine and Health, Chinese Academy of Sciences, were used for this study. All the animal experiments in this study were conducted in accordance with the Animal Protection Guidelines of Guangzhou Institutes of Biomedicine and Health, Chinese Academy of Sciences, Guangzhou, China.

### CRISPR plasmids for gene targeting

The pX330 vector carrying Cas9 expression cassette and blank gRNA bone was obtained from Addgene. sgRNAs for gene targeting at *ROSA26* locus in HEK293 genome, *TP53*, *LMNA*, and *POU5F1* loci in porcine genome were designed on the website http://www.rgenome.net (Park et al., 2015). The primers were synthesized by Genewiz. sgRNA sequences were cloned into BpiI-digested pX330 plasmids by primer pairs annealing and T4 DNA ligase. Detailed sgRNA sequences were shown in **Supplemental Table 6C**.

### Design and Construction of ReCOIN cassette and targeting donors

The ReCOIN cassette carrying a pair of FRT sites, a pair of wild type LoxP sites, a pair of LoxP2272 sites, an inverted EGFP followed by rßglpA, a puromycin resistance selection cassette, and the highly conserved 5’ and 3’ splice sites. Detailed sequences were shown in Figure 1-figure supplement 1. The cassette was synthesized by Guangzhou IGE Biotechnology.

The pCMV-mCherry-pA expression plasmid with ReCOIN cassette inserted in the middle of mCherry was constructed. Briefly, mCherry-L, mCherry-R and ReCOIN cassette were PCR amplified and cloned into NheI (Thermo Fisher) and NotI (Thermo Fisher) digested pCMV-mCherry expression plasmid using pEASY-UniSeamless Cloning and Assembly Kit (TransGen Biotech) according to manufacturer’s instruction. Then, the pCMV-Cas9-pA expression plasmid with ReCOIN cassette inserted in the middle of Cas9 cDNA was constructed using the same method.

For knocking ReCOIN cassette into any desired genomic loci, the universal intermediate vector pFlexibleDT-ReCOIN was constructed through recombination of ReCOIN cassette into NotI (Thermo Fisher) and HindIII (Thermo Fisher)-digested pFlexibleDT using ClonExpress MultiS One Step Cloning Kit (Vazyme Biotech). Subsequently, hRosa26-Cas9-ReCOIN, p*LMNA*-ReCOIN, p*TP53*-ReCOIN, and p*POU5F1*-ReCOIN targeting donors described above were generated based on the universal intermediate vector pFlexibleDT-ReCOIN. The p*LMNA*-ReCOIN targeting vector containing 1042 bp 5′ homologous arm and 1008 bp 3′ homologous arm, and the p*POU5F1*-ReCOIN targeting vector containing 1218 bp 5′ homologous arm and 1004 bp 3′ homologous arm were constructed. Briefly, left and right homologous arms around the ReCOIN cassette insertion sites, including *ROSA26* locus in HEK293 genome, *TP53*, *LMNA*, and *POU5F1* loci in porcine genome, were amplified by the KOD One PCR Master Mix (Toyobo) and purified by the HiPure Gel Pure DNA Mini Kit (Magen). And then the left and right homologous arms were successively inserted into linearized pFlexibleDT-ReCOIN using ClonExpress MultiS One Step Cloning Kit (Vazyme).

In addition, to achieve drug-inducible Cre and Flp expression, the targeting vector comprising of a TRE3G driving Cre and Flp-ERT2 expression cassette, a blasticidin resistance selection cassette, and left and right homologous arms at Hipp11 locus was constructed by inserting above elements into NotI (Thermo Fisher) and HindIII (Thermo Fisher)-linerized pFlexibleDT using pEASY-UniSeamless Cloning and Assembly Kit (TransGen Biotech).

### Human embryonic kidney (HEK) 293 cell culture and plasmid transfection

HEK 293 cells were cultured in medium consisting of high glucose DMEM (Hyclone) supplemented with 10% fetal bovine serum (Gibco) and 1× penicillin-streptomycin (Gibco), and they were maintained in a cell incubator at 37°C and 5% CO_2_. Cell cryopreservation solution consisted of 10% DMSO (Sigma) and 90% FBS (Gibco). We used a Neon transfection system (Life Technology), at the condition of 1150V, 20ms, and 2 pulses, with 100µL reaction volume of resuspension (R) buffer (Thermo Fisher), mixed respectively with CRISPR plasmids and targeting donor plasmids, expression plasmids of pCMV-mCherry inserted with ReCOIN allele, Cre, or Flp, for gene targeting or transient expression in HEK 293 cells negative for mycoplasma. In brief, for transient expression in HEK 293 cells, 5 µg expression plasmids of pCMV-mCherry and pCMV-Cas9 inserted with ReCOIN allele, Cre, or Flp were electroporated into 100 thousand cells, respectively. For gene targeting and generating of ReCOIN-Cas9 HEK 293 cell line, 5 µg pX330 plasmids containing sgRNA for human *ROSA26* locus and 5 µg Cas9-ReCOIN donors were electroporated into 100 thousand cells. One day post electroporation, HEK 293 cells were dissociated and seeded into 6 cm culture dishes. And 1 µg/mL puromycin was added to the culture medium to remove negative cells. PCR reactions to identify the genotype of these single-cell-derived colonies were conducted, with the primers listed in Supplemental Table 6A.

### Porcine fetal fibroblast (PFF) culture and plasmid transfection

PFFs derived from the 35-day-old Bama miniature pig fetuses were isolated and cryopreserved as previous study described (Wu et al., 2018). We used the culture medium of PFF consisting of high glucose DMEM (Hyclone) supplemented with 15% fetal bovine serum (Gibco), 1×Non-Essential Amino Acids (Gibco), 1×GlutaMax (Gibco), 1×sodium pyruvate (Gibco), 1× penicillin-streptomycin (Gibco) and 1 µg/mL L-ascorbic acid (Sigma). PFFs were maintained in a cell incubator at 38.5°C and 5% CO_2_. For gene targeting in PFFs, PFFs were thawed and seeded into 10 cm plate the day before electroporation. The next day, the PFFs reached an approximate 80% confluency. We used the Neon transfection system (Life Technology), at the condition of 1350V, 30ms, and 1 pulse, with 100-µL reaction volume of resuspension (R) buffer (Thermo Fisher), mixed with CRISPR plasmids and targeting donor plasmids for electroporation in these PFFs. In brief, for screening of ReCOIN-P53 PFFs, 10 µg pX330 plasmids containing gRNA for *TP53* and 10 µg ApaLI (Thermo Fisher)-linerized TP53-ReCOIN targeting donors were electroporated into about 1 million PFFs. After a 24-hour recovery, the electroporated PFFs were digested with 0.25% trypsin and suspended with PFF culture medium. Then these cells were counted and equally divided into twenty 10 cm culture dishes with 5000 cells each. The culture medium was refreshed and 300 ng/µL puromycin were added to maintain and select positive cells every other day. We picked up the single-cell-derived colonies by cloning cyclinders (Corning) after about 8-10 days. After 3-6 days growth of these cells, 90% were passaged, and 10% were lysed and identified by 5′- and 3′-junction fragment PCR analysis using primers listed in Supplemental Table 6A. Positive cell colonies with correct gene knock-in were expanded and cryopreserved for further use. For screening of CIRKO-TP53, CIRKO-POU5F1, and CIRKO-LMNA, 10 µg pX330 plasmids containing gRNA for *TP53 or LMNA or POU5F1*, 10 µg ApaLI (Thermo Fisher)-linerized targeting donors, 10 µg pX330 plasmids containing gRNA for *HIPP11*, 10 µg ApaLI (Thermo Fisher)-linerized *HIPP11* targeting donors described above, were electroporated into 2 million PFFs containing ROSA26-rtTA elements as our previous study reported. The procedures for picking up single-cell-derived colonies, subculture and genotype identification were the same as above description. Notely, 3 µg/µL blasticidin and 300 ng/µL puromycin were added to select positive cells in turns.

### High-throughput DNA sequencing and data analysis

We used amplification primers containing Illumina forward and reverse adapters (Supplemental Table 6B) for a first round of PCR amplifying the genomic regions containing the gRNA targeting sites. In Brief, we performed 10-µL PCR reactions with 0.5 µM of each forward and reverse primer, 1 µL of extracted genomic DNAs and 5 µL of 2× Hot Start High Fidelity Q5 Master Mix (NEB). PCR reactions were carried out under the following condition: 95 °C for 3 min, then 30 cycles of [98 °C for 30 s, 61 °C for 20 s, and 72 °C for 30 s], followed by a final 72 °C extension for 5min. Then, we added unique Illumina barcoding primer pairs to different samples in a secondary round of PCR reaction. Specifically, 50-µL PCR reactions containing 0.5 µM of each unique forward and reverse Illumina barcoding primer pair, 2 µL of unpurified the first round of PCR reaction mixture, and 25 µL of 2× Hot Start High Fidelity Q5 Master Mix (NEB) were performed, under the following condition: 95 °C for 3 min, then 12 cycles of [98 °C for 30s, 61 °C for 20 s, and 72 °C for 30 s], followed by a final 72 °C extension for 5 min. Purified products of the secondary round PCR reaction were subjected to Annoroad Gene Technology Corporation (Beijing) for next-generation sequencing. Finally, we used a previously reported tool named CRISPResso2 (Clement et al., 2019) to analyze the high-throughput DNA sequencing data.

### T7EN1 cleavage assay

First, we performed PCR to amplify the DNA sequences around target sites using the 2 × Rapid Taq Master Mix (Vazyme). Then, we purified the PCR products using the HiPure Gel Pure DNA Mini Kit (Magen). Approximate 200 ng PCR product of each reaction was annealed and incubated with T7EN1 (Vazyme) in accordance with the manufacture’s instruction. At last, we quenched the reactions by adding 1.5 µL of 0.25 M EDTA, and the T7EN1-digested PCR products were analyzed with 2% agarose gel.

### Flow cytometry analysis

For only analyzing the percentage of EGFP or mcherry expression in HEK 293 cells or PFFs, cell samples were prepared by using the 0.05% or 0.25% trypsin to dissociate adherent cells, 1×PBS to wash and resuspend cells and the 100 mesh strainer to remove cell debris. Then the prepared samples were analyzed by using an Accuri C6 flow cytometer (Accuri Cytometers). For analyzing and collecting the EGFP positive PFFs, the prepared samples were subjected to a MoFlo Astrios (BD).

### RNA extraction and RT-PCR

Total RNA was extracted using RNeasy Mini kit (Qiagen) with an on-column DNase digestion (Qiagen). We performed reverse transcription reactions by using approximate 500 ng of RNA as template, in a 10 µL reaction volume with the PrimeScript RT Reagent Kit with gDNA Eraser (Takara) following the recommended protocol. RT-PCR was conducted by using the diluted products of reverse transcription reactions as template and using 2 × Rapid Taq Master Mix (Vazyme) according to the manufacturer’s protocol, with the primers listed in Supplemental Table 6A. And the products of RT-PCR were analyzed by using 1% agarose gel electrophoresis and were subjected to Sanger sequencing.

### RNA-sequencing and data analysis

The harvested ReCOIN-Cas9 HEK 293 cells were stored in RNAlater™ Stabilization Solution (Thermo Fisher, AM7020) and then cells were subjected to Annoroad Gene Technology Corporation (Beijing) for library construction and transcriptome sequencing. We performed quality control for the obtained data using FastQC (version v0.11.9). Then we removed unqualified and adapter sequences by using fastp, with parameters “--detect_adapter_for_pe”. We used human genome from NCBI (GCF_000001405.39) as the reference and mapped remained reads through accessible gene annotation provided by Hisat2 (version 2.1.0) with default parameters. We processed BAM files by using Samtools (version 1.9). And then we used Htseq-count (version 0.11.2) to quantify the gene expression levels. Next, we applied custom scripts to generate gene expression matrices. We performed differential expression analysis of protein-coding genes by using the R/Bioconductor package DESeq2 (version 1.30.0), with standard parameters. Finally, we conducted KEGG pathway and Gene Ontology analyses based on differentially expressed genes through clusterProfiler (version 3.18.0).

### Generation of ReCOIN-TP53 pigs by somatic cell nuclear transfer

Screened ReCOIN-P53 PFFs were thawed and seeded in 24-well plate before SCNT. Cells that reached a state of contact inhibition were preferentially used as nucleus donors. SCNT protocol was conducted as previously described (Lai and Prather, 2003). *In vitro* matured oocytes were enucleated by gently aspirating the first polar body and adjacent cytoplasm with glass pipette in manipulation medium supplemented with 6.7 µg/mL cytochalasin B (Sigma, C6762). The donor cell was subsequently injected into the perivitelline space, followed by fusion of the donor cell with the enucleated oocyte and activation by electrical pulses (1.2 kV/cm, 30 ms, 2 pulses). The reconstructed embryos were then transferred to embryo culture medium and cultured in a cell incubator at 38.5°C and 5% CO_2_ for 20 hours. The day after the estrus was observed, reconstructed embryos were transferred into the oviducts of wild type surrogates by surgery. After embryo transfer for 30-35 days, an ultrasound scanner was used to determine the pregnancy status. Pregnant sows were assessed by periodic monitoring to track the development of cloned embryos. The cloned piglets carrying ReCOIN-P53 element were delivered by receipt sows natural birth after about 114 days of gestion.

### Genomic DNA extraction and genotyping

Genomic DNAs were collected from one small piece of ear tissues of cloned newborn piglets using TIANamp Genomic DNA Kit (TIANGEN). Then the genomic DNAs were used as a PCR template and PCR products were analyzed by agarose gel electrophoresis for identification of the cloned piglet genotypes. The primers used for PCR were the same as those used for cell-level genotyping.

### Cell immunofluorescence staining

ReCOIN-P53 and CIRKO-P53 PFFs were seeded onto 1% gelatin-coated 24 well plates. Then cells were fixed in 4% paraformaldehyde (PFA) for 15 minutes at room temperature and 1×PBS was used to wash cells thrice for 5 minutes each time. Then cells were permeabilized with 0.5% triton X-100 (Sigma) and 10% goat serum (Panera) in PBS for 30 minutes. Next, cells were incubated with primary antibodies P53 (1:200, Proteintech) and EGFP (1:500, Proteintech) overnight at 4 °C. On the following day, the cells were washed thrice with 1×PBS for 5 min each time. Then cells were incubated with secondary antibodies (1:1000, Cell Signaling Technology) for 2 h at room temperature. And then the cells were washed thrice with PBS for 5 min each time. Finally the cells were mounted with sealing agent containing DAPI (Sigma). Fluorescent images were captured with an inverted fluorescent microscope (Olympus).

### CIRKO-pPOU5F1 PFFs for SCNT

Screened CIRKO-pPOU5F1 PFFs were treated with/out Dox (1 µg/mL) and 4-OHT (2.5 µM) before used as donor cells for SCNT, which was performed as described above. The reconstructed embryos were cultured *in vitro* for 6 days to blastocyst stage, when single blastocyst was collected into PCR tube with 10 µL lysis buffer for subsequent PCR reactions. Also, a part of blastocysts were subjected to fixing with PFA, followed by immunofluorescence staining. Briefly, the blastocysts were fixed with 4% PFA at room temperature for 30 min, permeabilized with 2% Triton X-100 (Sigma) and 5% goat serum (Panera) in PBS for 30 min, and blocked with 5% goat serum for 60 min, sequentially. Next, the embryos were incubated with primary antibodies against POU5F1 (1:200, Santa Cruz) and EGFP (1:500, Proteintech) overnight at 4 °C. After washing with PBS, the embryos were incubated with appropriate secondary antibodies (1:1000, Cell Signaling Technology) for 1 h at room temperature. Finally, the nuclei were stained with sealing agent containing DAPI (Sigma), and fluorescent images were captured with an laser scanning confocal microscopy (Zeiss).

### Statistical analysis

In this study, experiments were not randomized. Experiment arrangements were not blinded to the investigators during statistical and assessment process. We used Prism 7 (GraphPad) to perform the statistical analysis. All results were presented as mean ± standard deviation, with more than three independent experiments. Unpaired Student’s *t*-test was used to calculate *p* values and *p* value < 0.05 was considered as statistically significant. Independent replicate experiments were performed and similar results were observed.

## Data Availability

The deep sequencing data reported in this paper is deposited in GEO, accession number GSE188595 (https://www.ncbi.nlm.nih.gov/geo/query/acc.cgi?acc=GSE188595).

## Author contributions

H.S., Q.J., F.C., K.W., and Liangxue L. designed this study. H.S., Q.J., and F.C. performed most of the experiments and analyzed the data. Z.O., C.L., and Q.Z. performed SCNT and generated cloned pigs. S.G. performed the bioinformatics analysis. X.L., Lei L., S.M., and Y.Y. provided technical assistance. H.S., K.W., and Liangxue L. prepared the manuscript. K.W. and Liangxue L. supervised this study. All authors read and approved the final manuscript.

## Funding

This work was financially supported by the National Key Research and Development Program of China Stem Cell and Translational Research (2021YFA0805903, 2017YFA0105103); the National Natural Science Foundation of China (32170542, 81941004, 32100410); the Key Research & Development Program of Hainan Province (ZDYF2021SHFZ052); Major Science and Technology Projects of Hainan Province (ZDKJ2021030); the 2020 Research Program of Sanya Yazhou Bay Science and Technology City (202002011); the Youth Innovation Promotion Association of the Chinese Academy of Sciences (2019347); the Young Elite Scientist Sponsorship Program by CAST (YESS20200024); the Biological Resources Progaramme, Chinese Academy of Sciences (KFJ-BRP-017-57); the Science and Technology Planning Project of Guangdong Province, China (2020B1212060052, 2021B1212040016, 2021A1515110838); the Science and Technology Program of Guangzhou, China (202002030382, 202007030003); the Research Unit of Generation of Large Animal Disease Models, Chinese Academy of Medical Sciences (2019-I2M-5-025).

## Declaration of interests

The authors declare that they have no conflict of interest.

---

**Figure 1-figure supplement 1.**
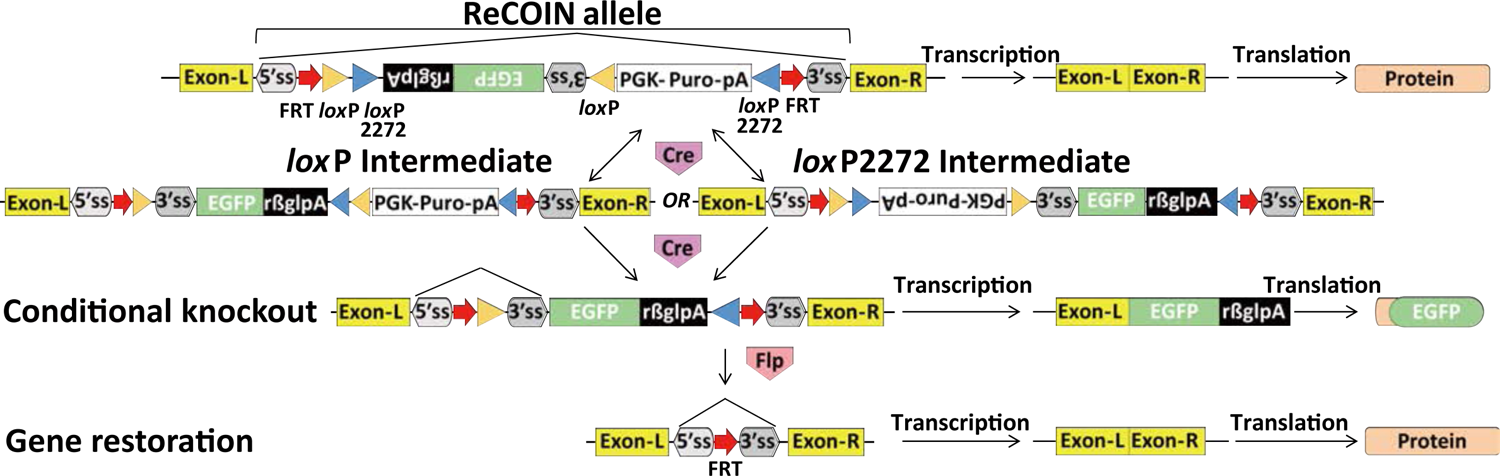
Detailed schematic of the ReCOIN cassette. Cre drives irreversible inversion of 3’ss (3’ splicing site), EGFP and the polyadenylation (PolyA) transcriptional terminator sequence, thereby terminating transcription of the target gene; Flpo drives deletion of 3’ss, EGFP and PolyA, thereby restoring gene’s expression.

**Figure 1-figure supplement 2.**
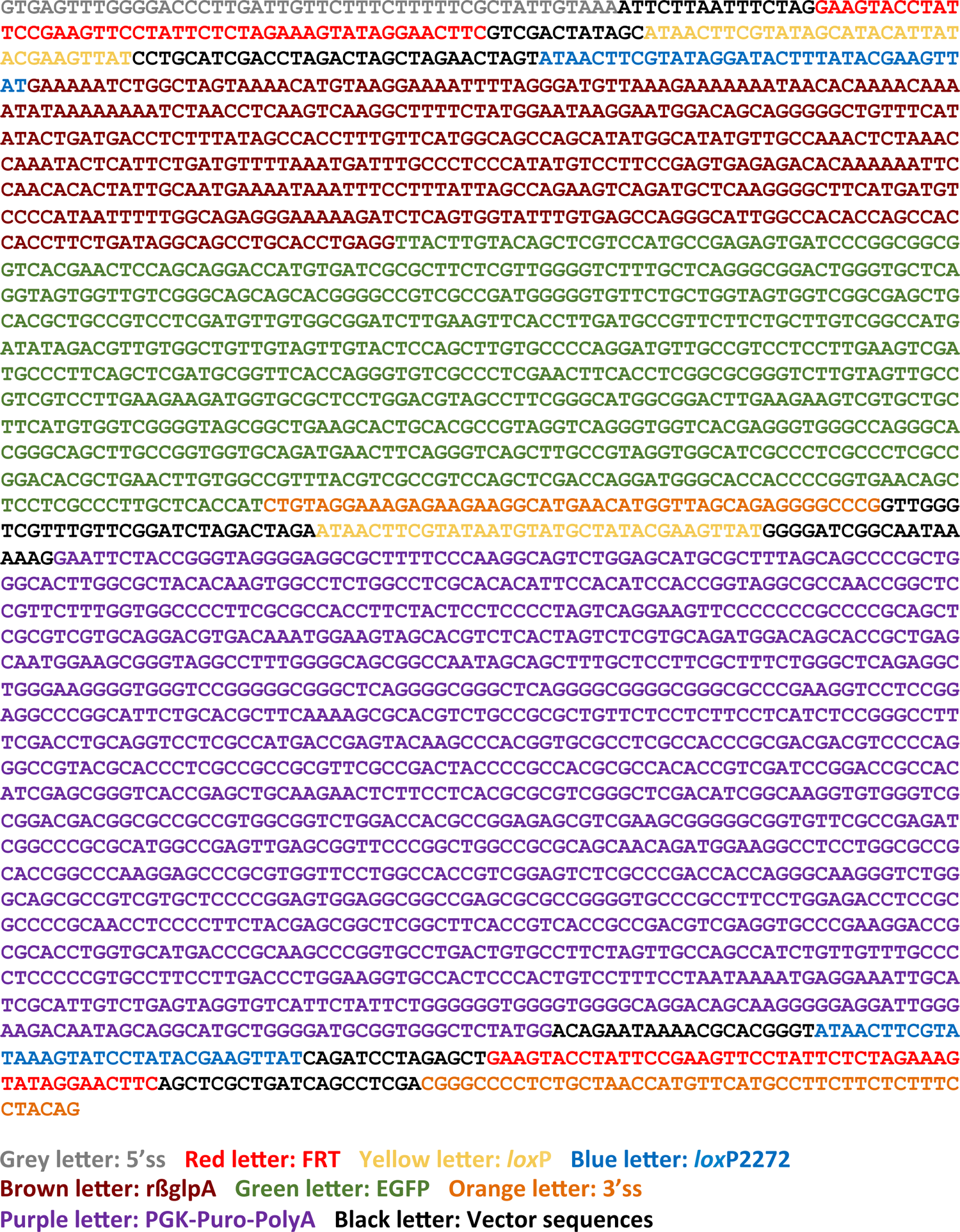
Nucleotide sequence for the ReCOIN construct.

**Figure 1-figure supplement 3.**
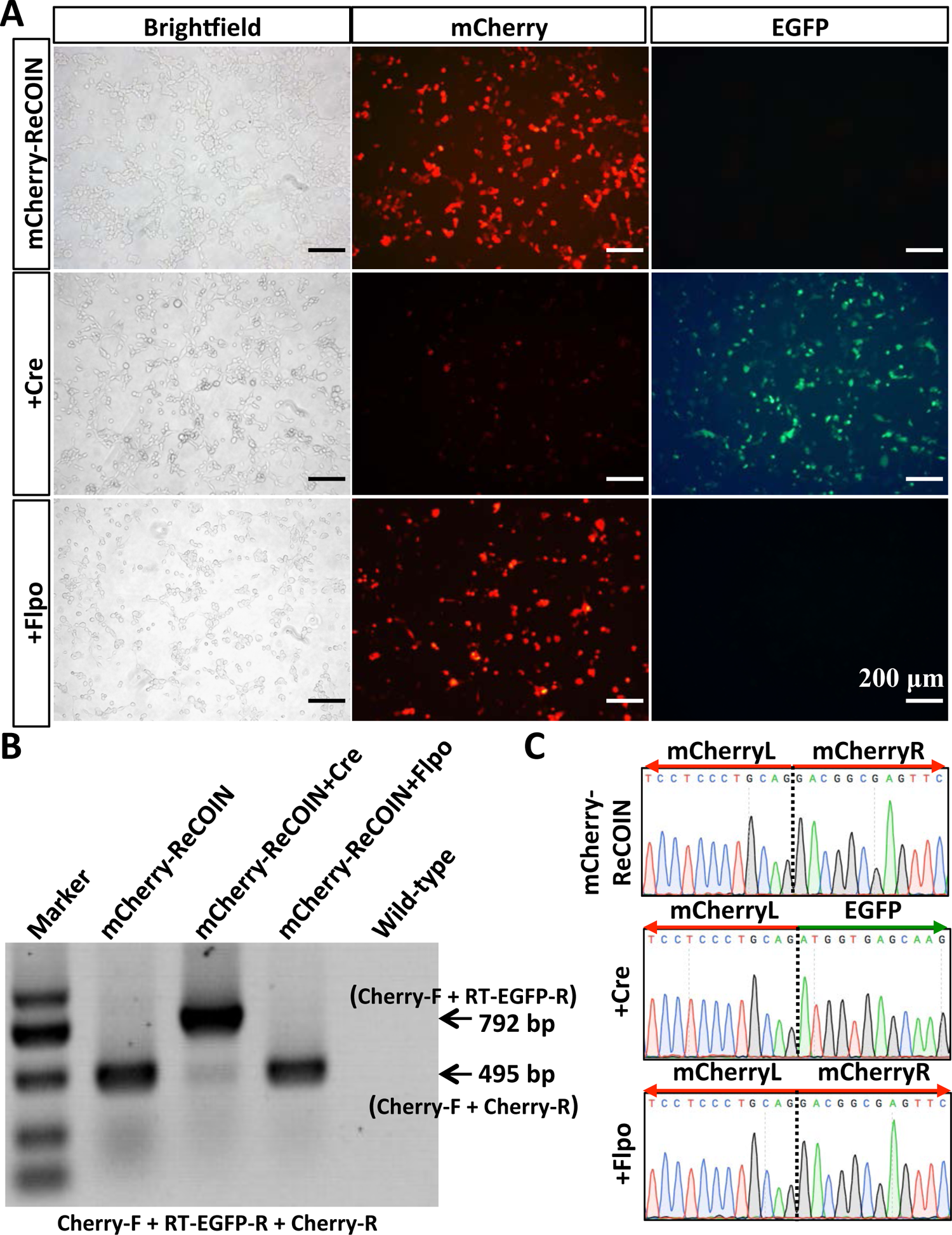
Validation of ReCOIN allele at mCherry by transient transfection with Cre and Flpo recombinase alone. (A) Fluorescent images of HEK293 cells transfected with the pCMV-mCherry-ReCOIN vector, pCMV-mCherry-ReCOIN vector+Cre-expressing vector, or pCMV-mCherry-ReCOIN vector+Flpo-expressing vector. mCherry protein is expressed in cells of the group transfected with pCMV-mCherry-ReCOIN alone (top row). Cre recombination disrupted mCherry and EGFP is expressed (middle row). Flpo recombination maintained the expression of mCherry (bottom row). (B) RT-PCR analysis of mCherry in HEK293 cells that were co-transfected with mCherry-ReCOIN and the indicated recombinase. All detected bands were of the expected size. (C) Sanger sequencing of RT-PCR products in (B).

**Figure 1-figure supplement 4.**
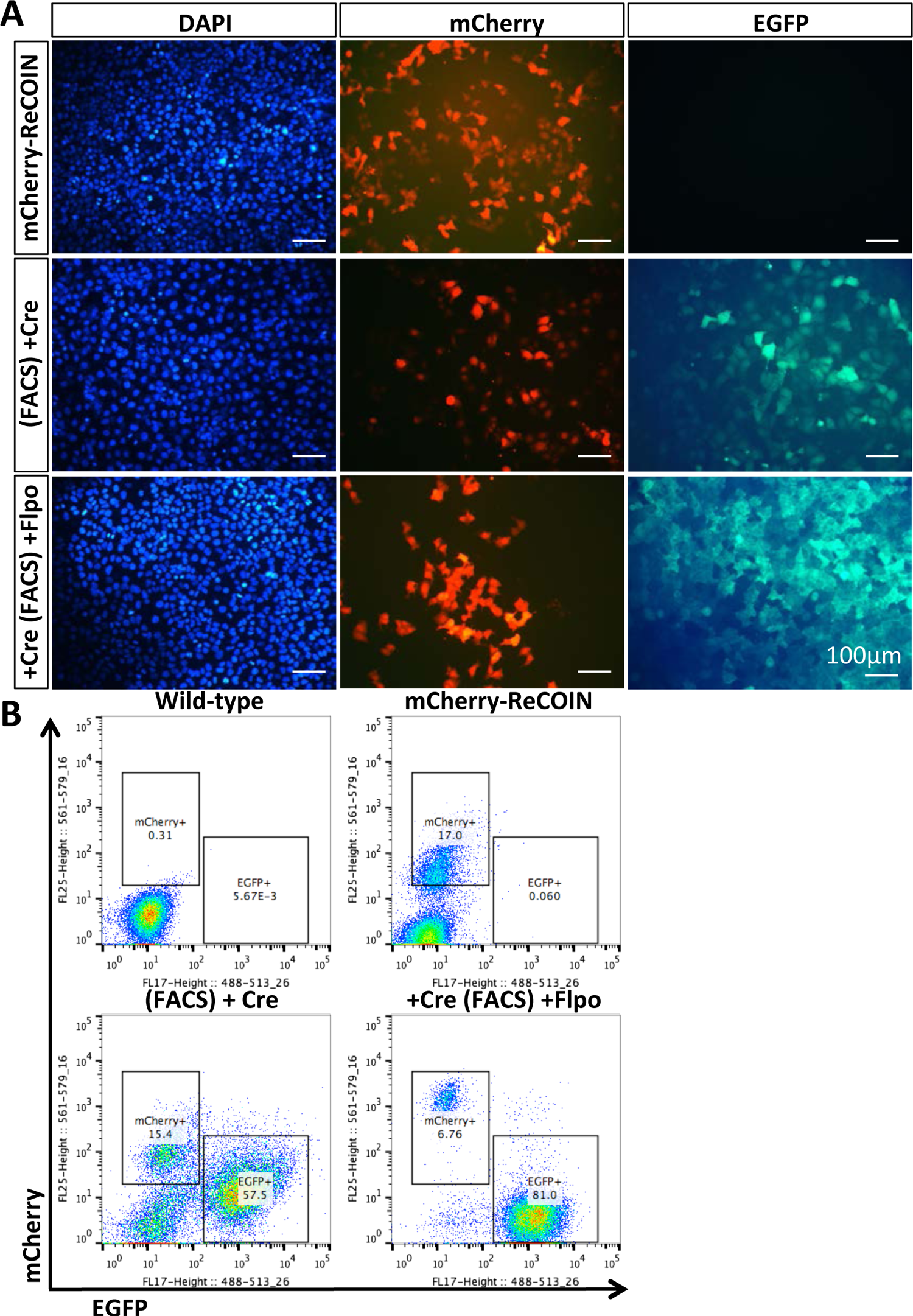
Validation of ReCOIN allele at mCherry by sequential transfection with Cre and Flpo recombinases. (A) Fluorescent images of the unsorted mCherry-ReCOIN cells, mCherry-ReCOIN+Cre cells, and mCherry-ReCOIN+Cre+Flpo cells. mCherry protein is expressed in the unsorted mCherry-ReCOIN cells (top row). Cre recombination disrupted mCherry and EGFP is expressed (middle row). Flpo recombination restored the expression of mCherry (bottom row). (B) Flow cytometry analysis of the unsorted mCherry-ReCOIN cells, mCherry-ReCOIN+Cre cells, and mCherry-ReCOIN+Cre+Flpo cells.

**Figure 1-figure supplement 5.**
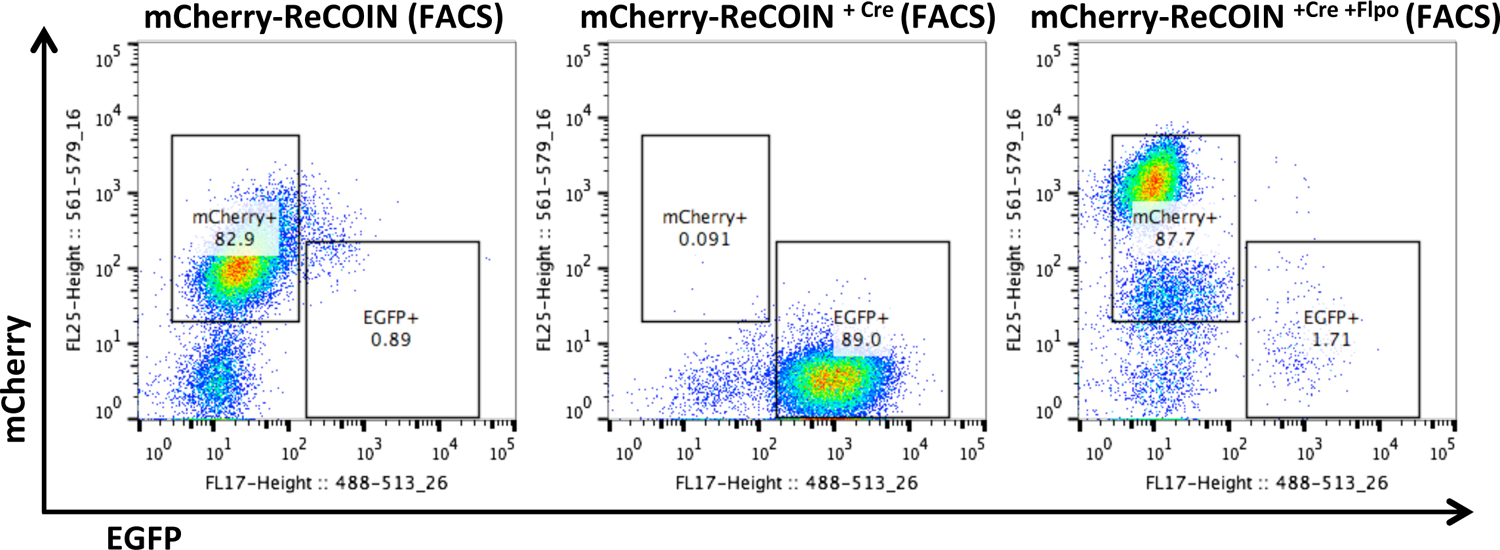
Flow cytometry analysis of the sorted mCherry-ReCOIN cells, mCherry-ReCOIN^+Cre^ cells, and mCherry-ReCOIN^+Cre+Flpo^ cells.

**Figure 1-figure supplement 6.**
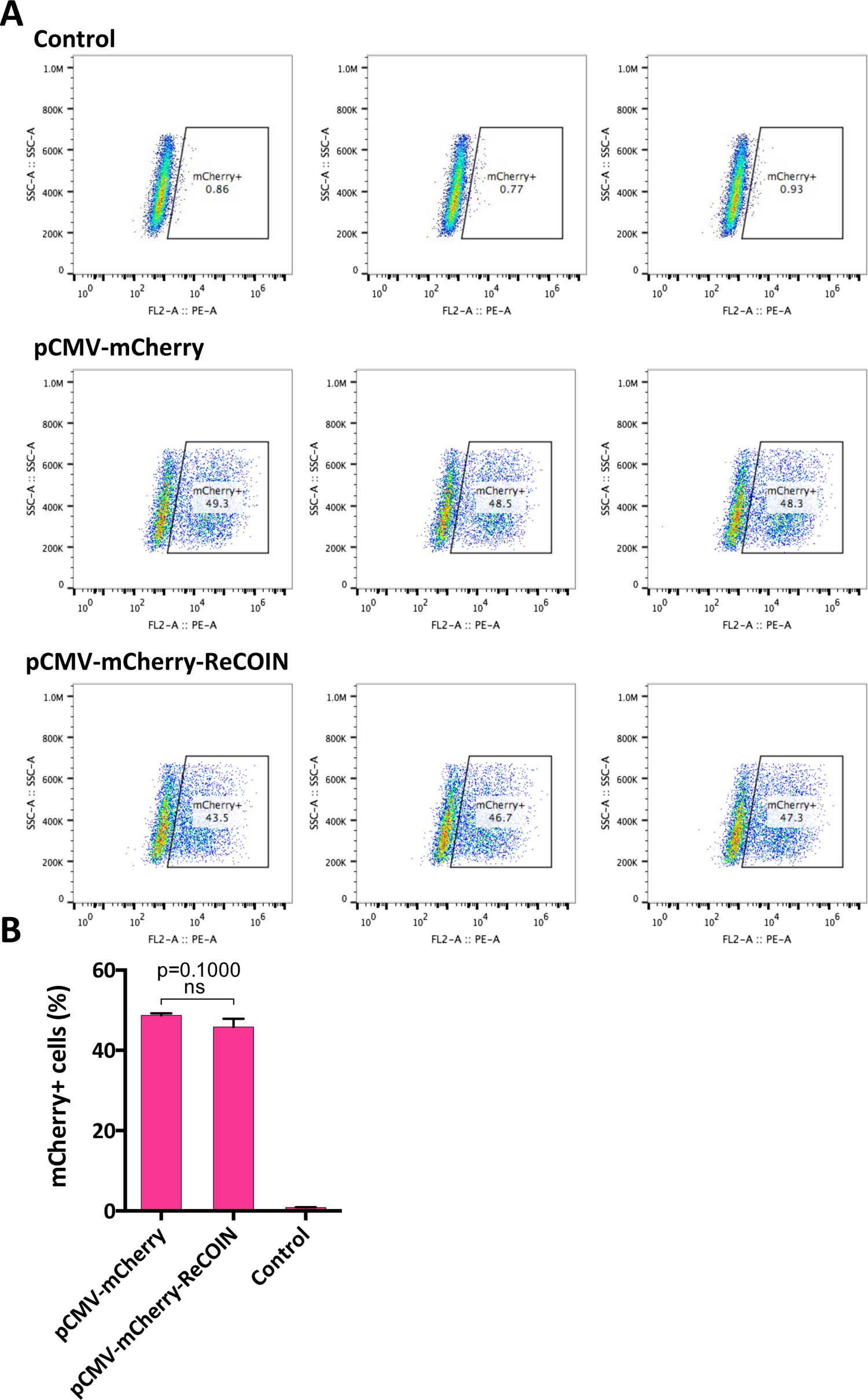
Evaluation of the expression level of mCherry in mCherry-ReCOIN. (A) Flow cytometry analysis of HEK293 cells transfected with the pCMV-mCherry or pCMV-mCherry-ReCOIN. (B) Bar plot of the flow cytometry results in (A). The data are presented as the mean ± SD of three independent experiments. Kruskal-Wallis test was used to calculate p values. ns indicates not significant.

**Figure 2-figure supplement 1.**
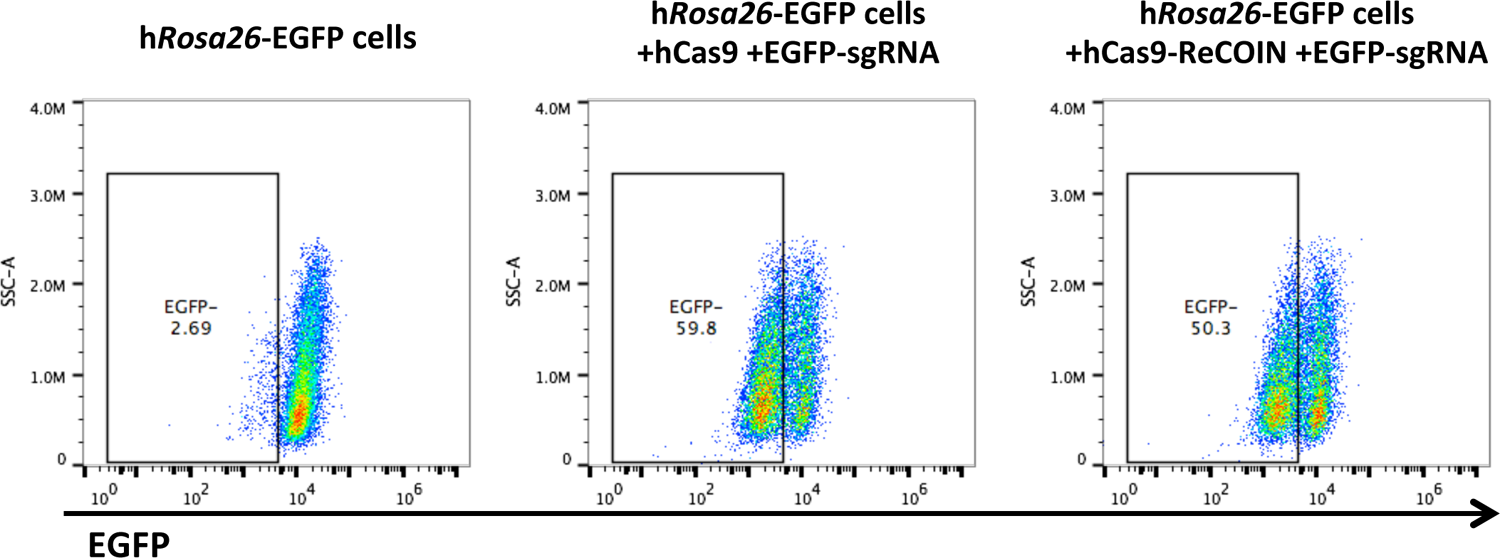
EGFP disruption assay using Cas9-ReCOIN and Cas9.

**Figure 2-figure supplement 2.**
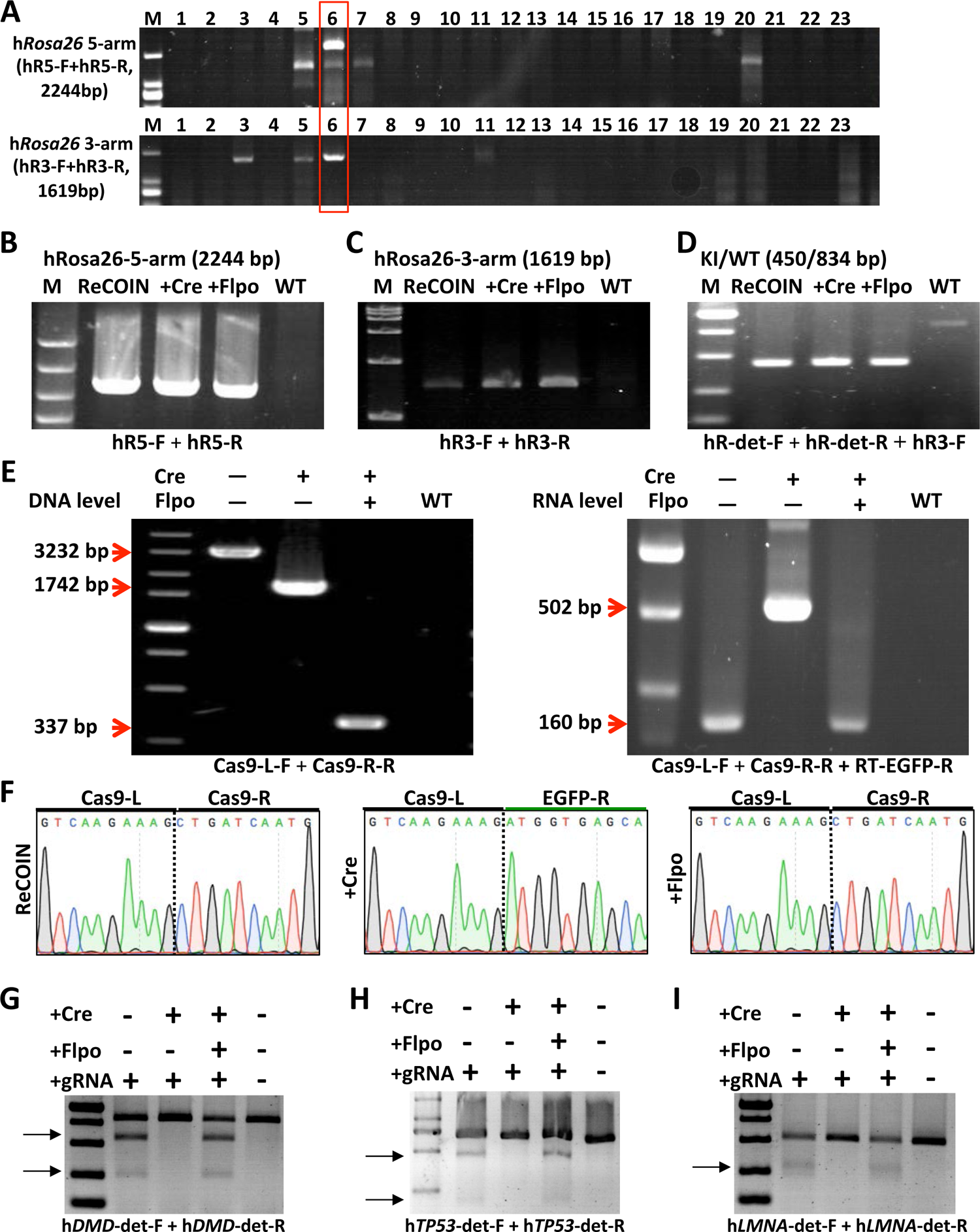
Insertion of the ReCOIN allele in the exogenous Cas9. (A) Detection of correctly integrated 5’ and 3’ arms by PCR in HEK293 colonies targeted with Cas9-ReCOIN. The primers used for genotyping are listed in Supplemental Table 6A. (B-D) PCR analysis confirmed the correct homologous recombination at the h*Rosa26* locus as detected by 5’ junction PCR (B), 3’ junction PCR (C) and homozygous/heterozygous PCR (D) in the indicated HEK293 colonies. All detected bands were of the expected size. (E) PCR-based (left column) and RT-PCR-based (right column) detection of ReCOIN allele within Cas9 in the indicated HEK293 colonies. (F) Sanger sequencing of the Cas9 cDNA in the indicated HEK293 colonies. (G-I) T7EN1-cleaved PCR amplicon fragments of the indicated HEK293 colonies transfected with DMD (G), TP53 (H) or LMNA (I) sgRNA.

**Figure 3-figure supplement 1.**
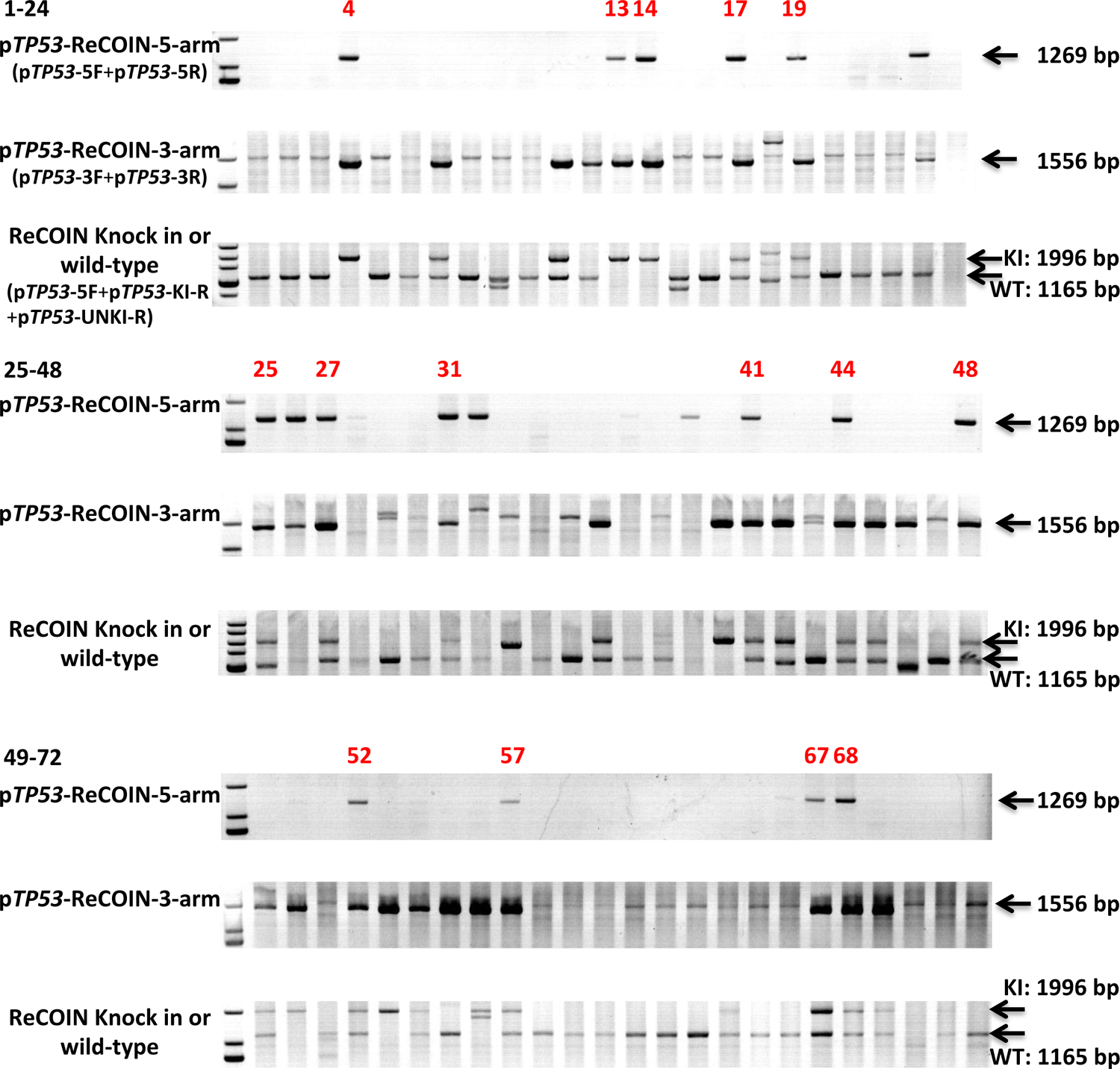
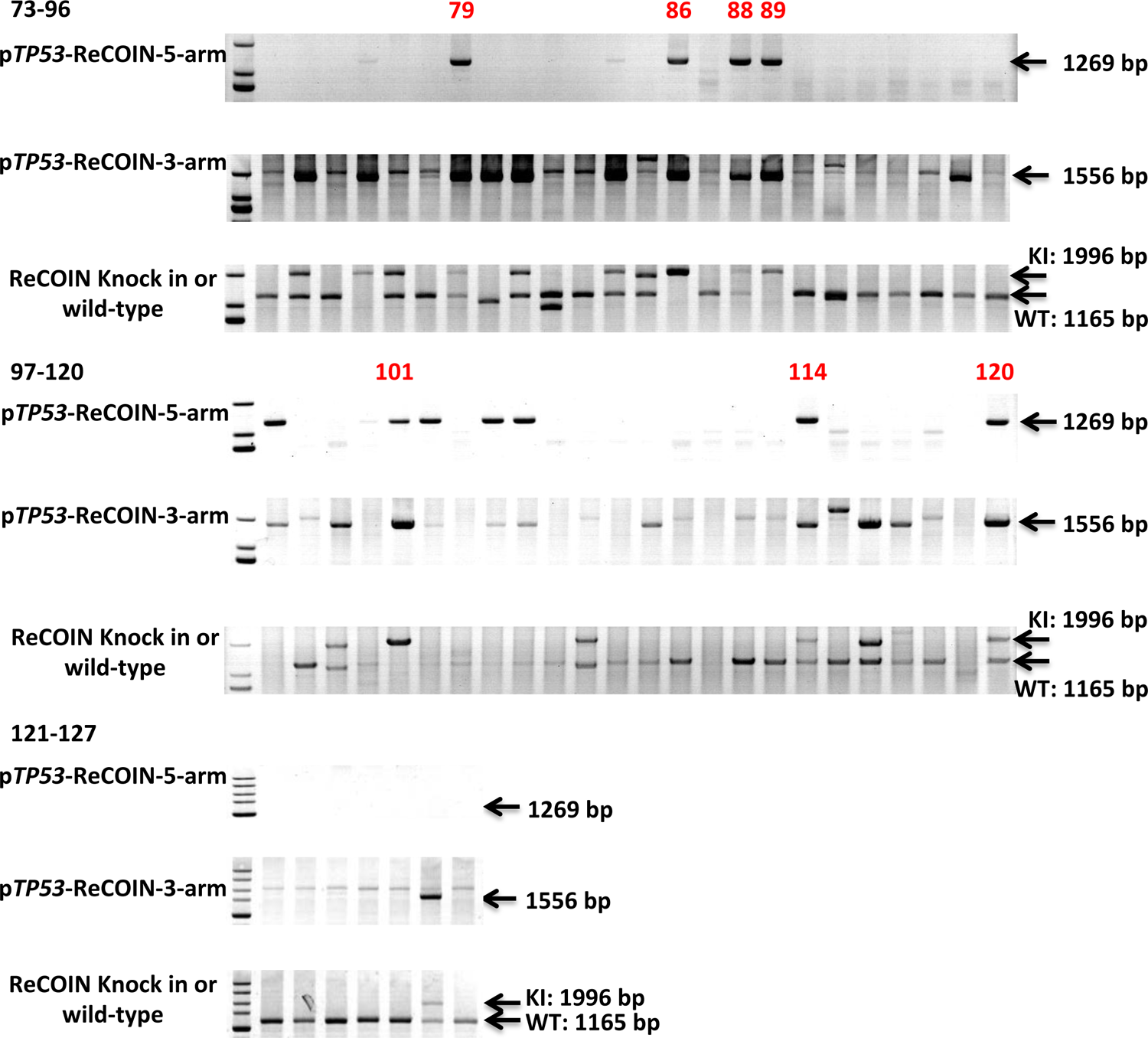
PCR analysis confirmed correct homologous recombination of the ReCOIN allele at p*TP53* locus in PFFs. Positive colonies are numbered in red.

**Figure 4-figure supplement 1.**
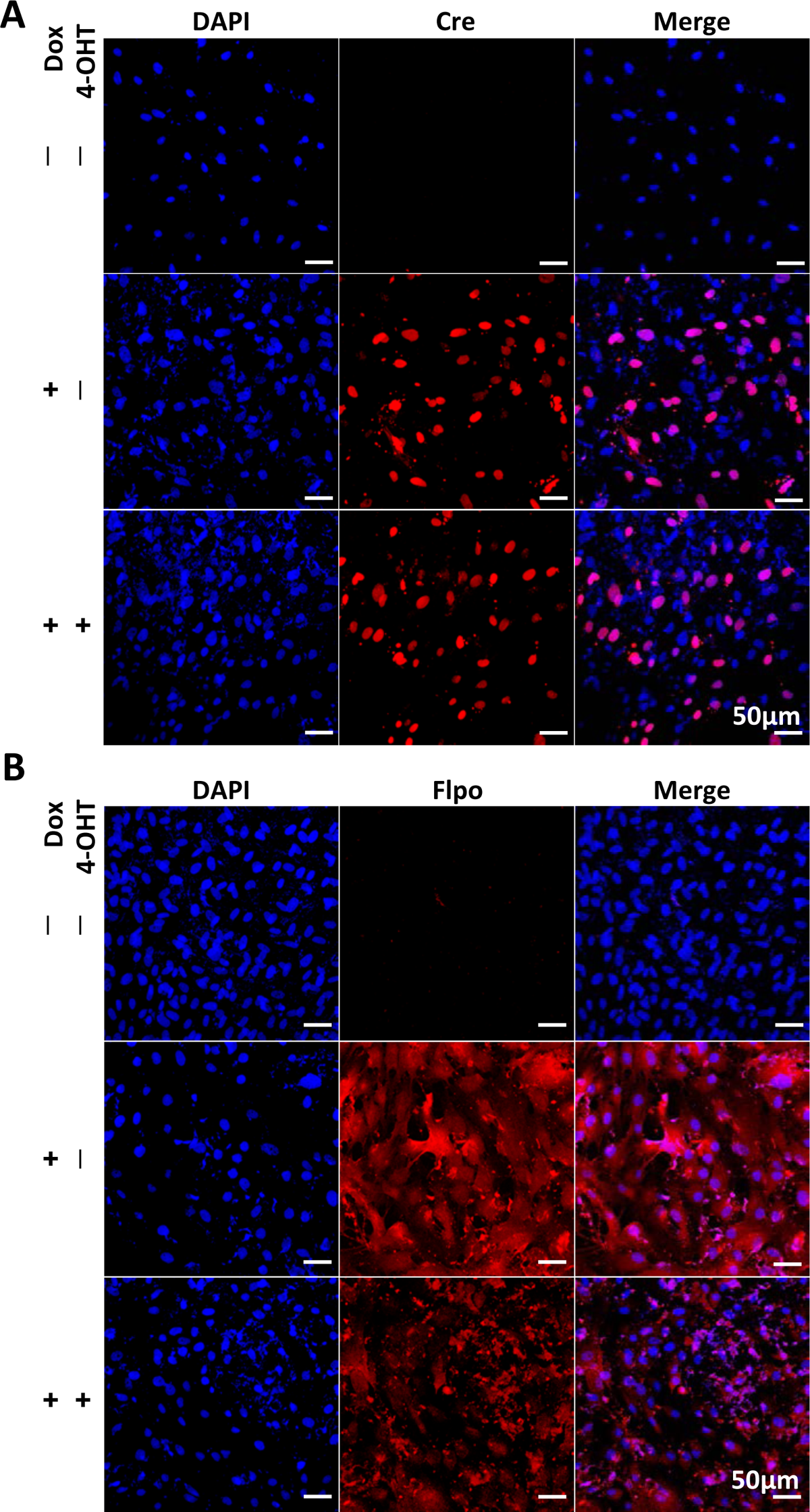
Immunocytochemical staining of Cre (A) and Flpo (B) in CIRKO PFFs at day 9 after treatment.

**Figure 5-figure supplement 1.**
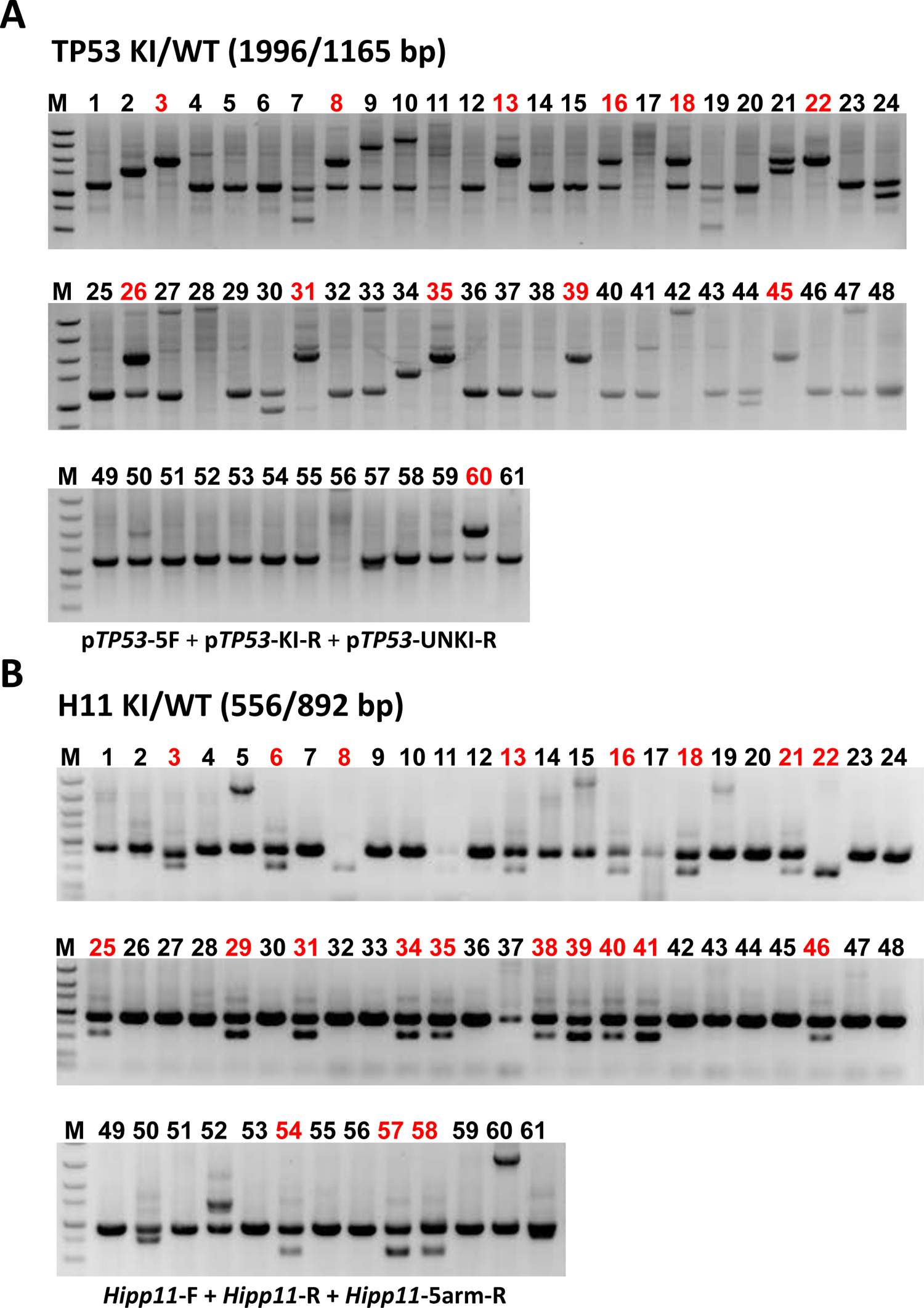
PCR analysis of CIRKO-p*TP53* colonies. (A) PCR analysis confirmed correct homologous recombination of the ReCOIN allele at p*TP53* locus in PFFs. Positive colonies are numbered in red. (B) PCR analysis confirmed correct homologous recombination of TRE3G-Cre-FlpERT2 at p*Hipp11* locus in PFFs. Positive colonies are numbered in red.

**Figure 5-figure supplement 2.**
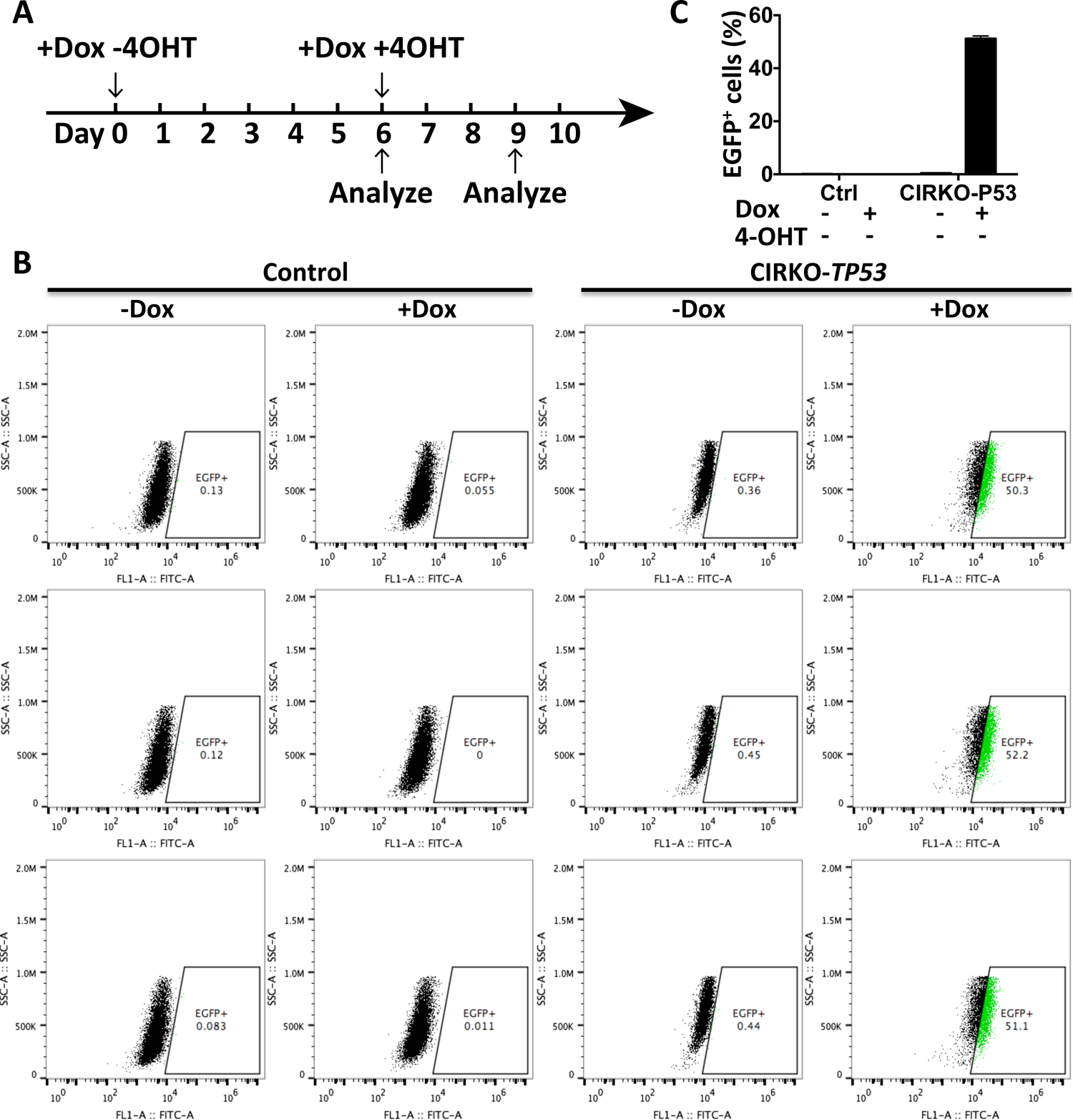
CIRKO-p*TP53* validates CIRKO system in PFFs at day 6 after treatment. (A) Flow chart for the treatment of CIRKO-p*TP53* PFFs. (B) Detection of EGFP reporter expression in the indicated CIRKO-p*TP53* PFFs by flow cytometry analysis at day 6 after treatment. Wild-type PFFs are used as controls. Triplicated biological experiments for the PFF clone were performed. (C) Histogram of EGFP-positive cell percentage in the indicated CIRKO-p*TP53* PFFs at day 6. Source data is available in Figure 5-figure supplement 2B.

**Figure 5-figure supplement 3.**
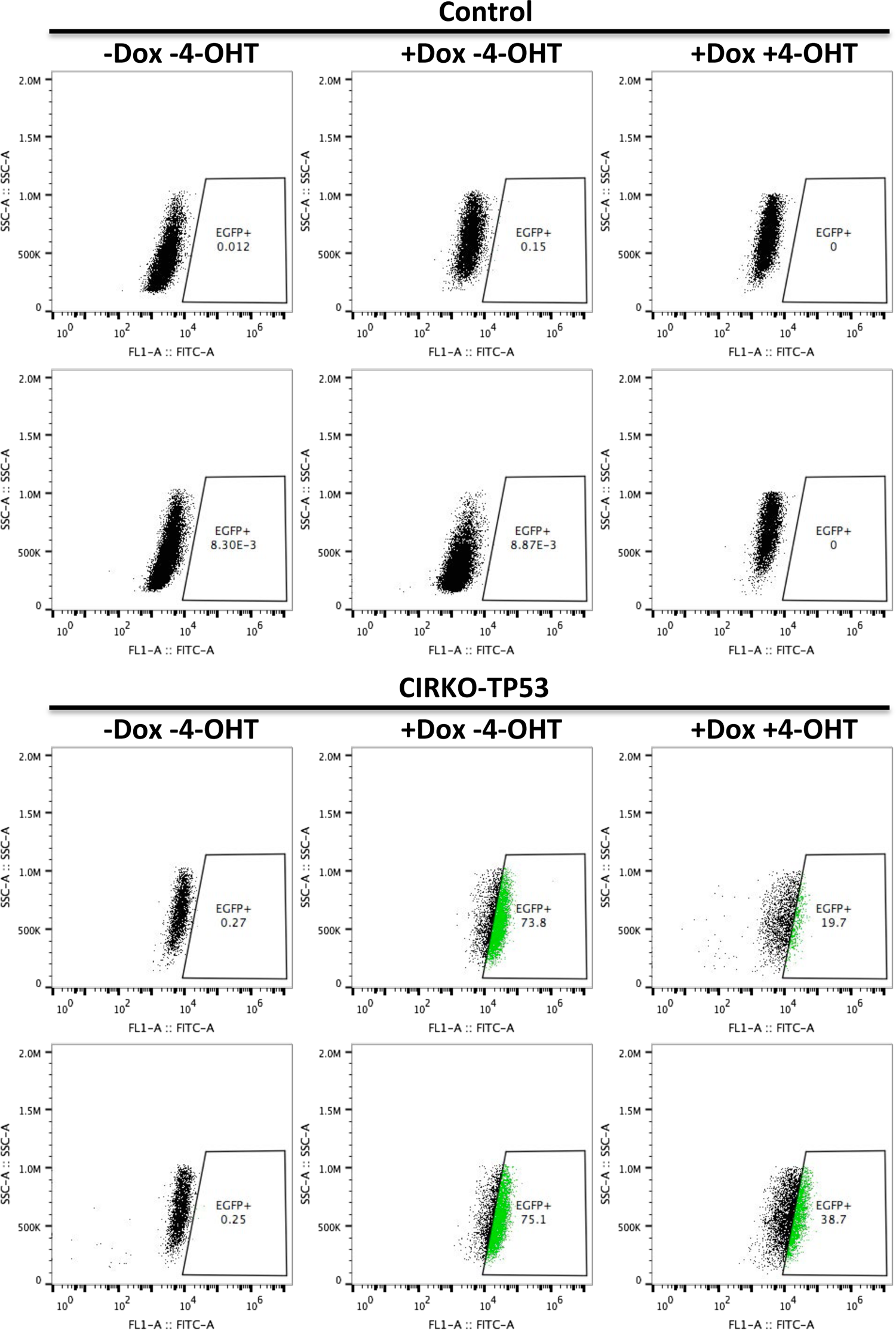
Detection of EGFP reporter expression in the indicated CIRKO-p*TP53* PFFs by flow cytometry analysis at day 9 after treatment. Wild-type PFFs are used as controls.

**Figure 5-figure supplement 4.**
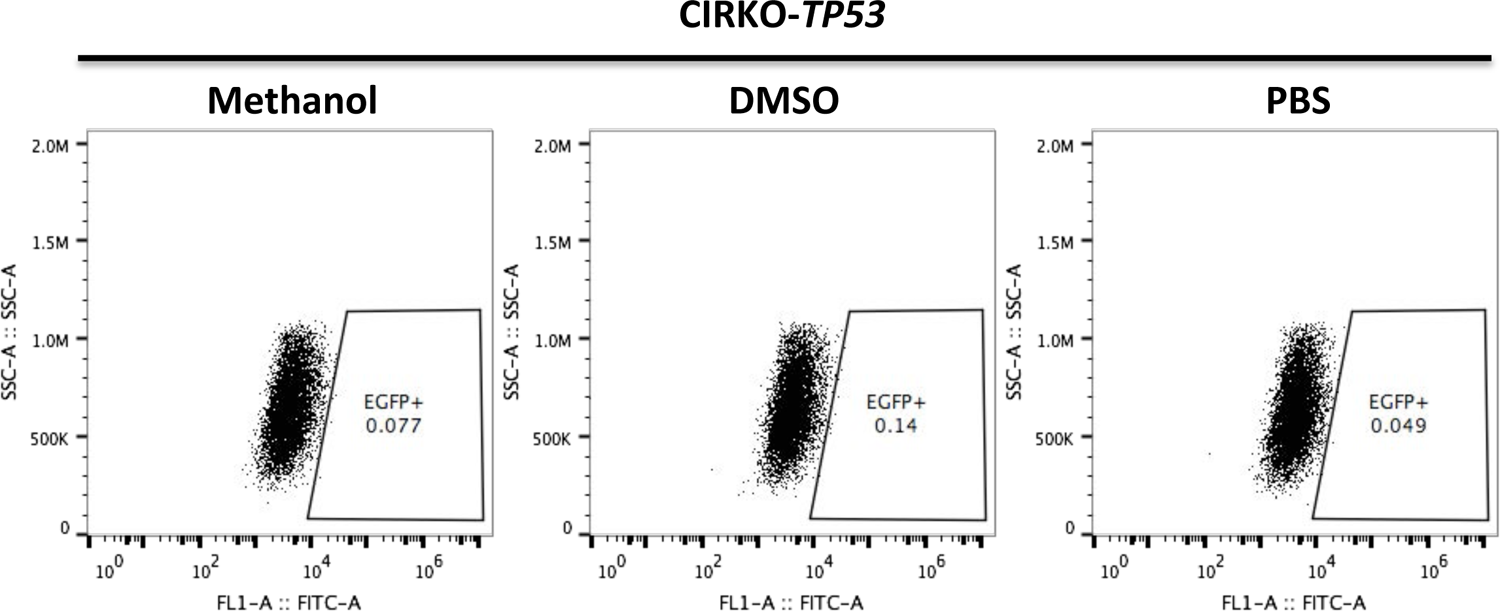
Detection of EGFP reporter expression in the indicated CIRKO-p*TP53* PFFs by flow cytometry analysis at day 9 after treatment.

**Figure 5-figure supplement 5.**
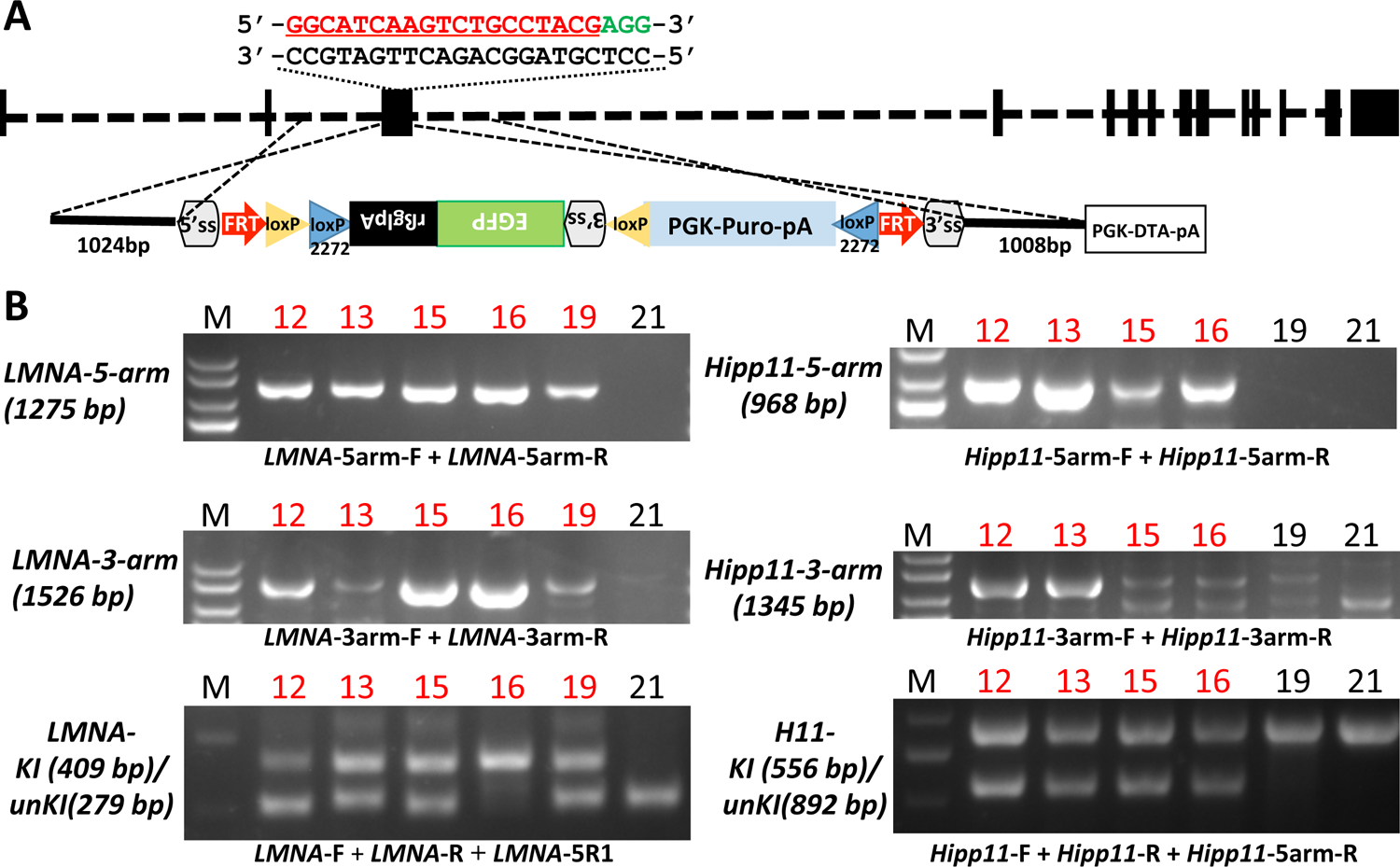
PCR analysis of CIRKO-p*LMNA* colonies. (A) Schematic diagram of the strategy for introducing ReCOIN at p*LMNA* via homologous recombination in PFFs. The sgRNA is used to target the third exon of p*LMNA* gene. The sgRNA guide sequence is indicated in red, and the PAM motif is indicated in green. (B) PCR analysis confirmed correct homologous recombination of the ReCOIN allele and TRE3G-Cre-FlpERT2 at p*LMNA* and p*Hipp11* locus, respectively. Positive colonies are numbered in red.

**Figure 6-figure supplement 1.**
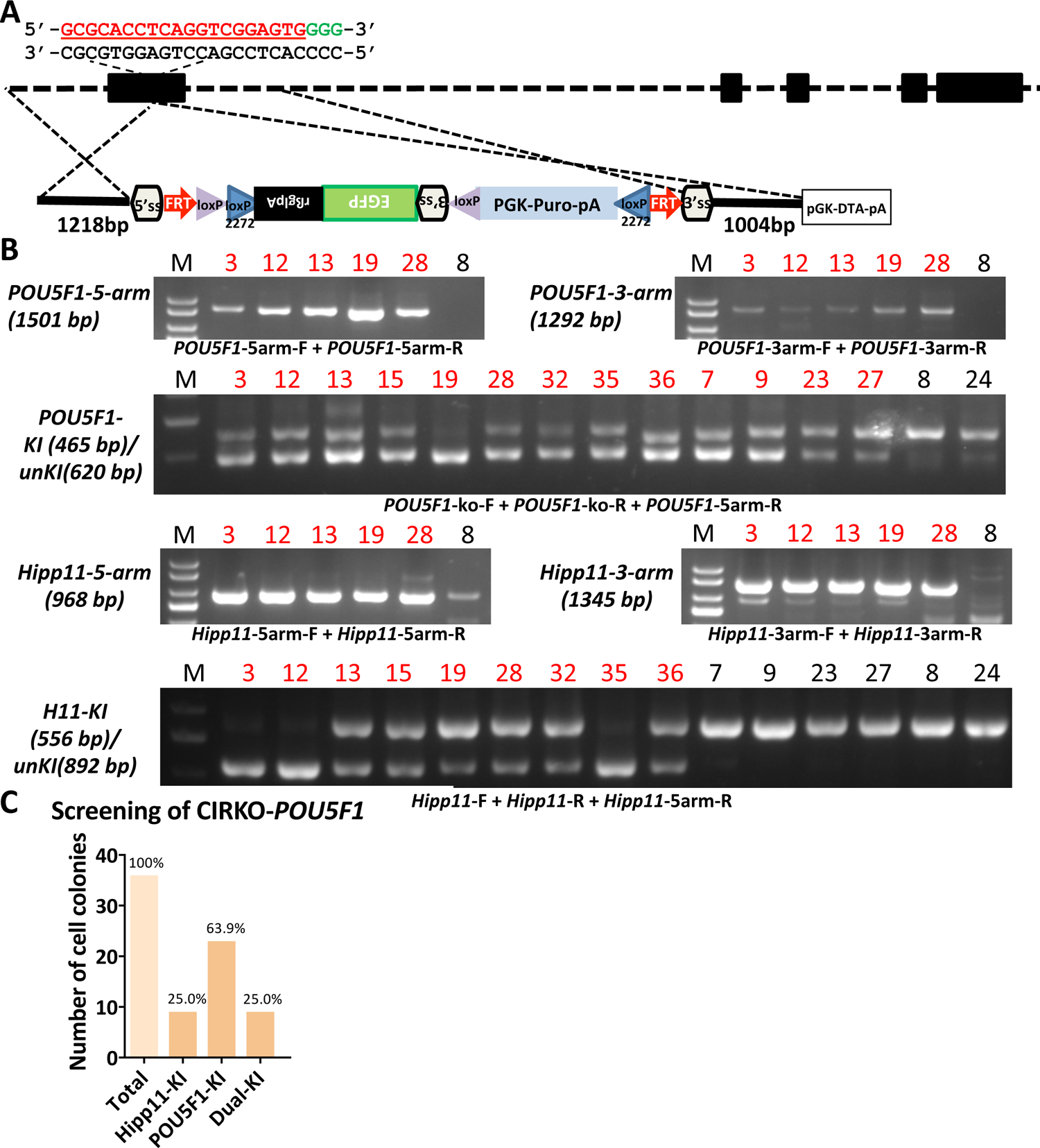
PCR analysis of CIRKO-p*POU5F1* colonies. (A) Schematic diagram of the strategy for introducing ReCOIN at p*POU5F1* via homologous recombination in PFFs. The sgRNA is used to target the first exon of p*POU5F1* gene. The sgRNA guide sequence is indicated in red, and the PAM motif is indicated in green. (B) PCR analysis confirmed correct homologous recombination of the ReCOIN allele and TRE3G-Cre-FlpERT2 at p*POU5F1* and p*Hipp11* locus, respectively. Positive colonies are numbered in red. (C) Histogram of screening CIRKO-p*POU5F1* colonies.

**Figure 6-figure supplement 2.**
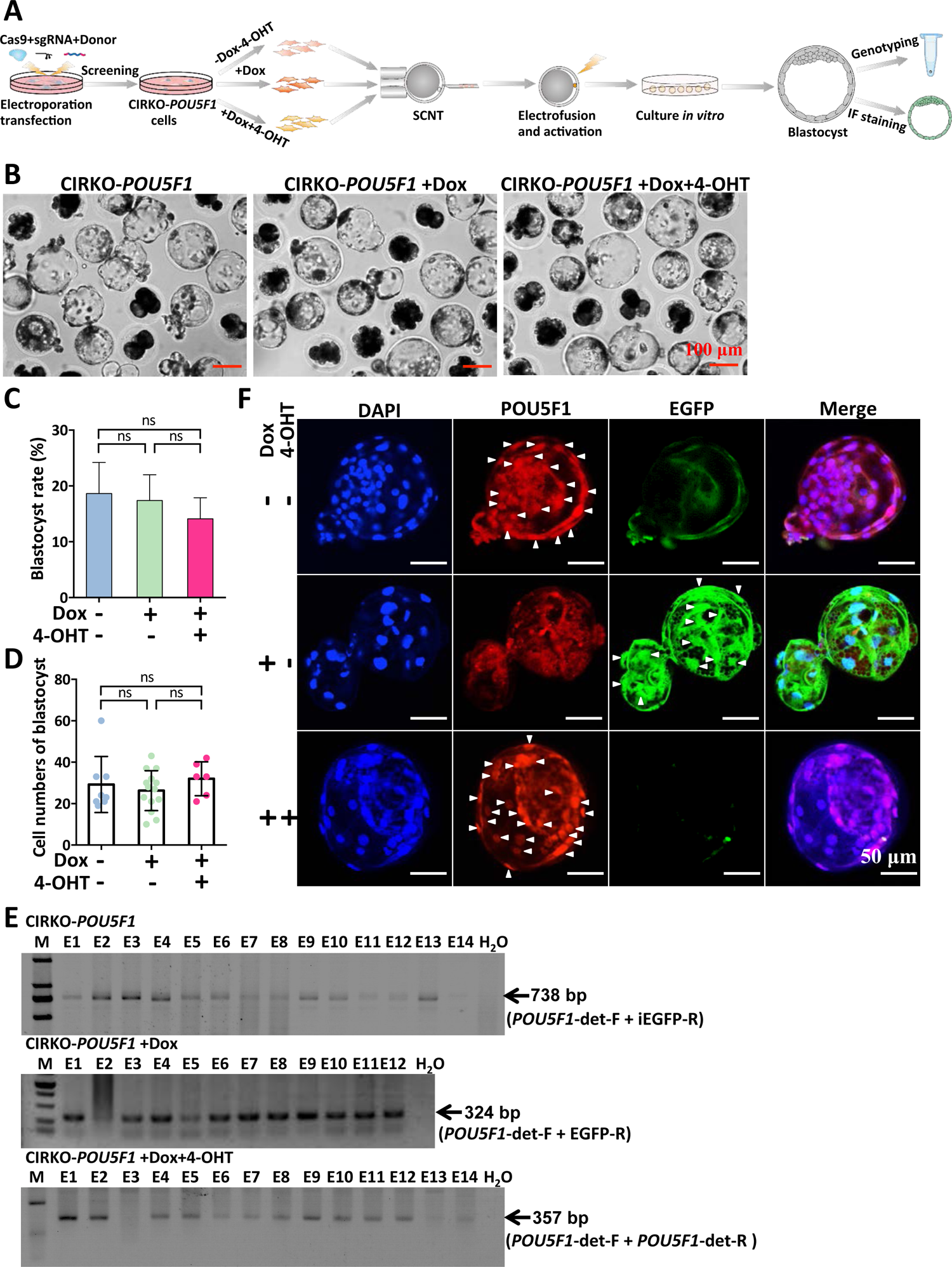
CIRKO facilitates reversible knockout of p*POU5F1* at the cellular level. (A) Experimental procedure to perform reversible knockout of p*POU5F1* at the cellular level. (B) Representative images of day 6 reconstructed embryos from untreated (left column), Dox-treated (middle column), and Dox+4-OHT treated (right column) CIRKO-p*POU5F1* PFFs. (C) Blastocyst rates of the reconstructed embryos from the indicated CIRKO-p*POU5F1* PFFs. The data are presented as the mean ± SD of three independent experiments. Kruskal-Wallis test was used to calculate p values. ns indicates not significant. Source data is available in Supplemental Table 4. (D) Total cell numbers of individual blastocyst from the indicated CIRKO-p*POU5F1* PFFs. The data are presented as the mean ± SD. Sample sizes were n=8, 14 and 6 for embryos from untreated, Dox-treated and Dox+4-OHT treated cells, respectively. Kruskal-Wallis test was used to calculate p values. ns indicates not significant. Source data is available in Supplemental Figure 16F and Supplemental Figure 17. (E) PCR analysis of p*POU5F1* locus in individual blastocyst from the indicated CIRKO-p*POU5F1* PFFs. (F) Representative confocal images of POU5F1 (red) and EGFP (green) expression in the blastocysts from the indicated CIRKO-p*POU5F1* PFFs. White arrows indicate overlap with nuclei.

**Figure 6-figure supplement 3.**
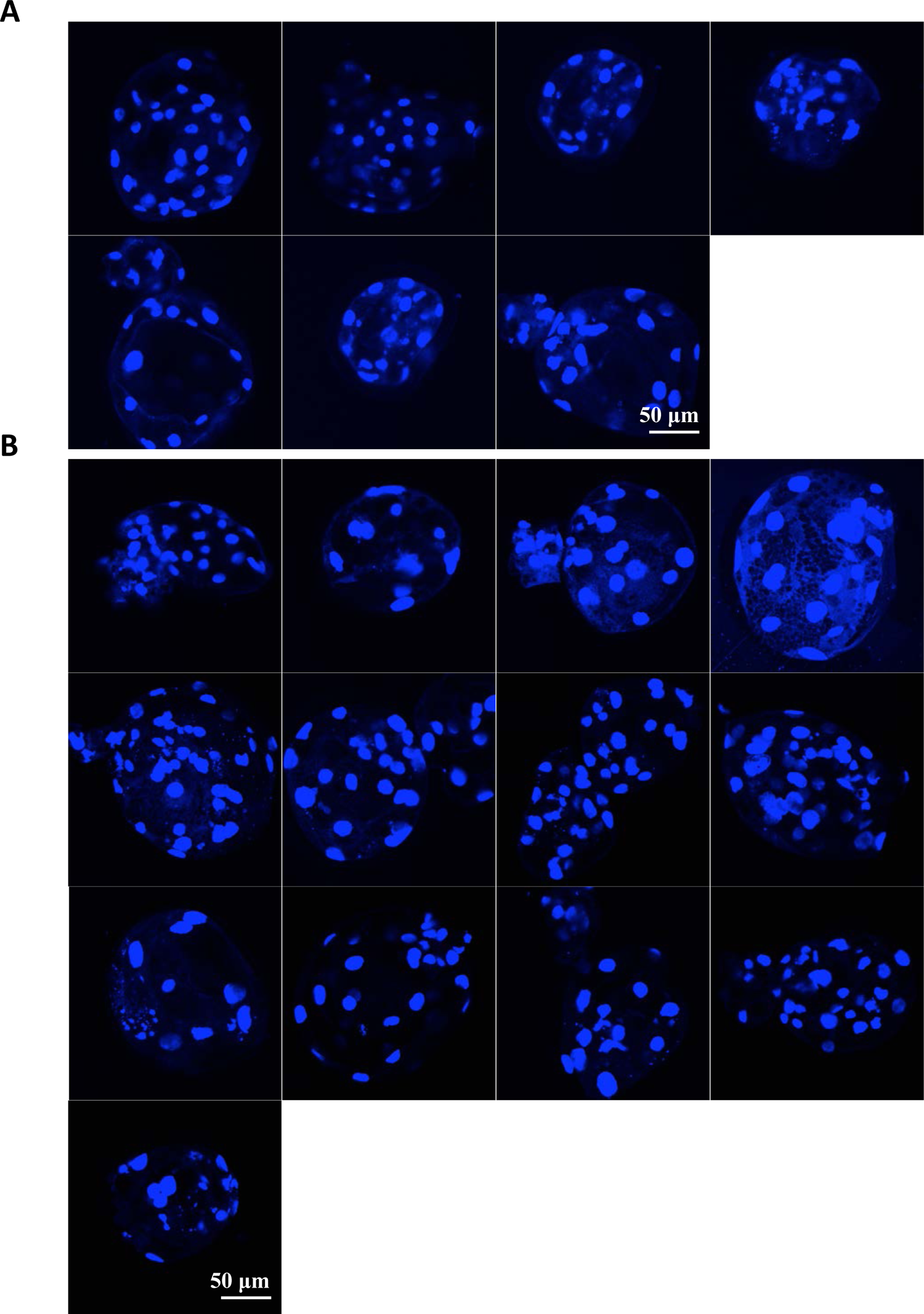
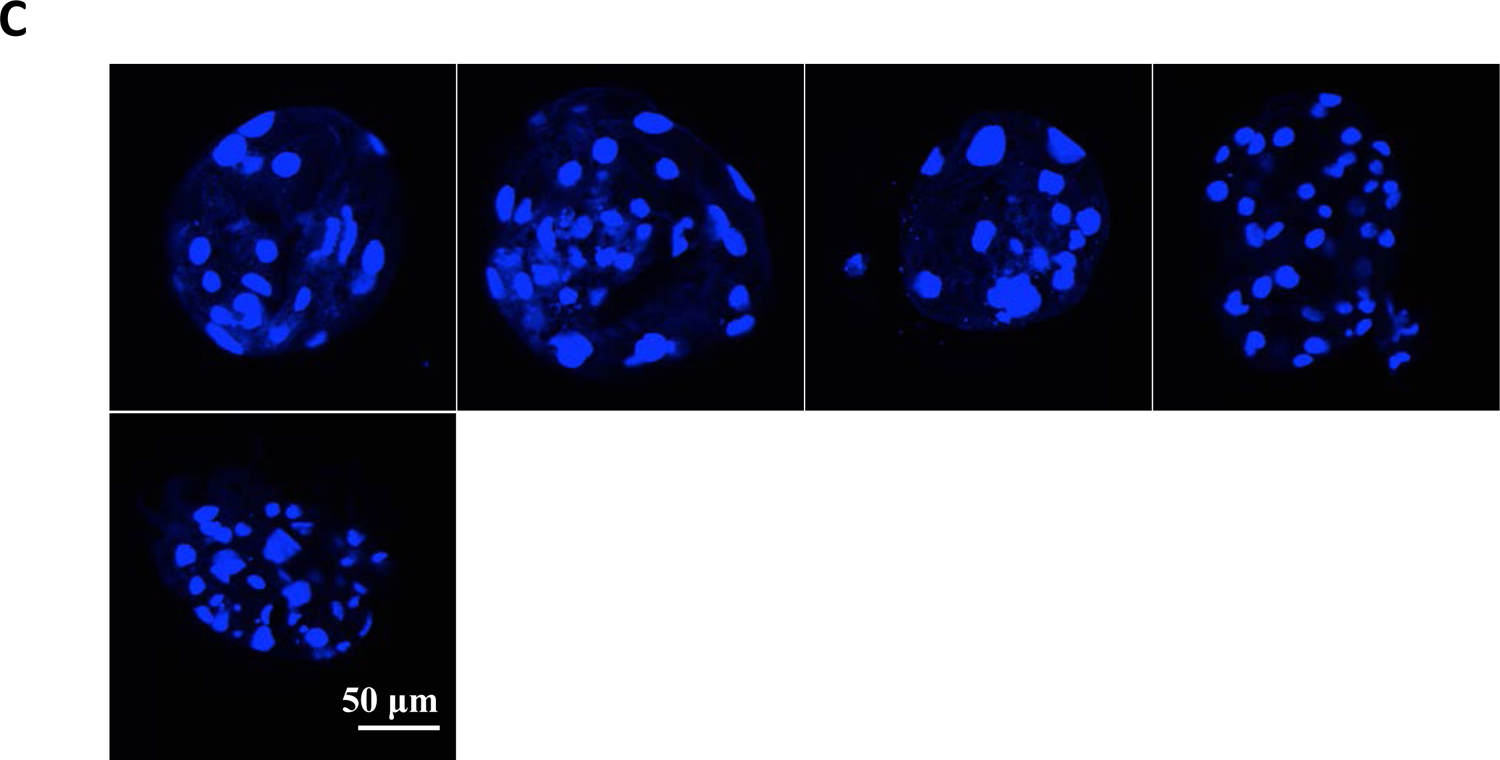
Confocal images for nucleus of individual blastocyst from untreated (A), Dox-treated (B), and Dox+4-OHT treated (C) CIRKO-p*POU5F1* PFFs.

**Figure 6-figure supplement 4.**
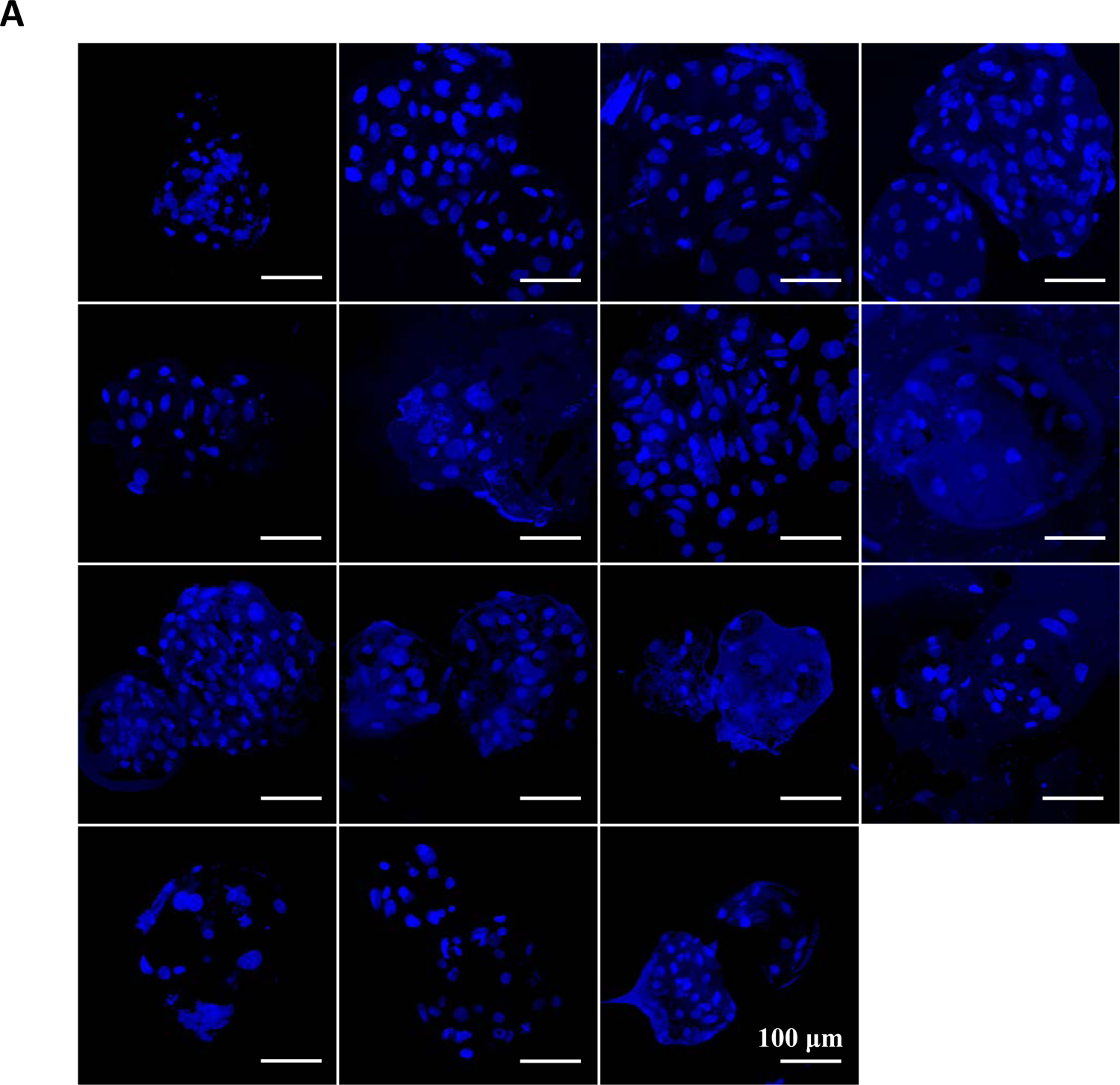
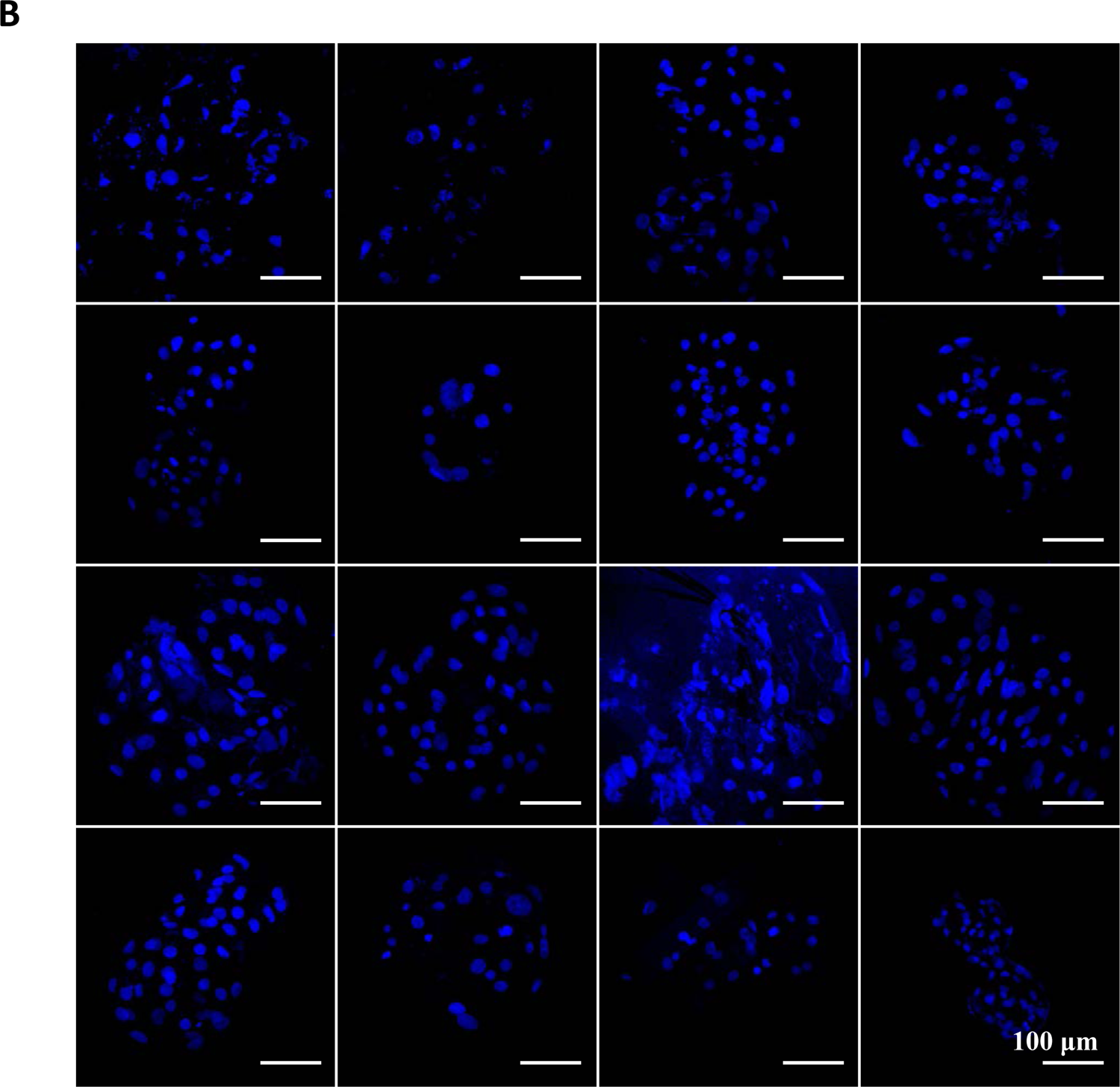
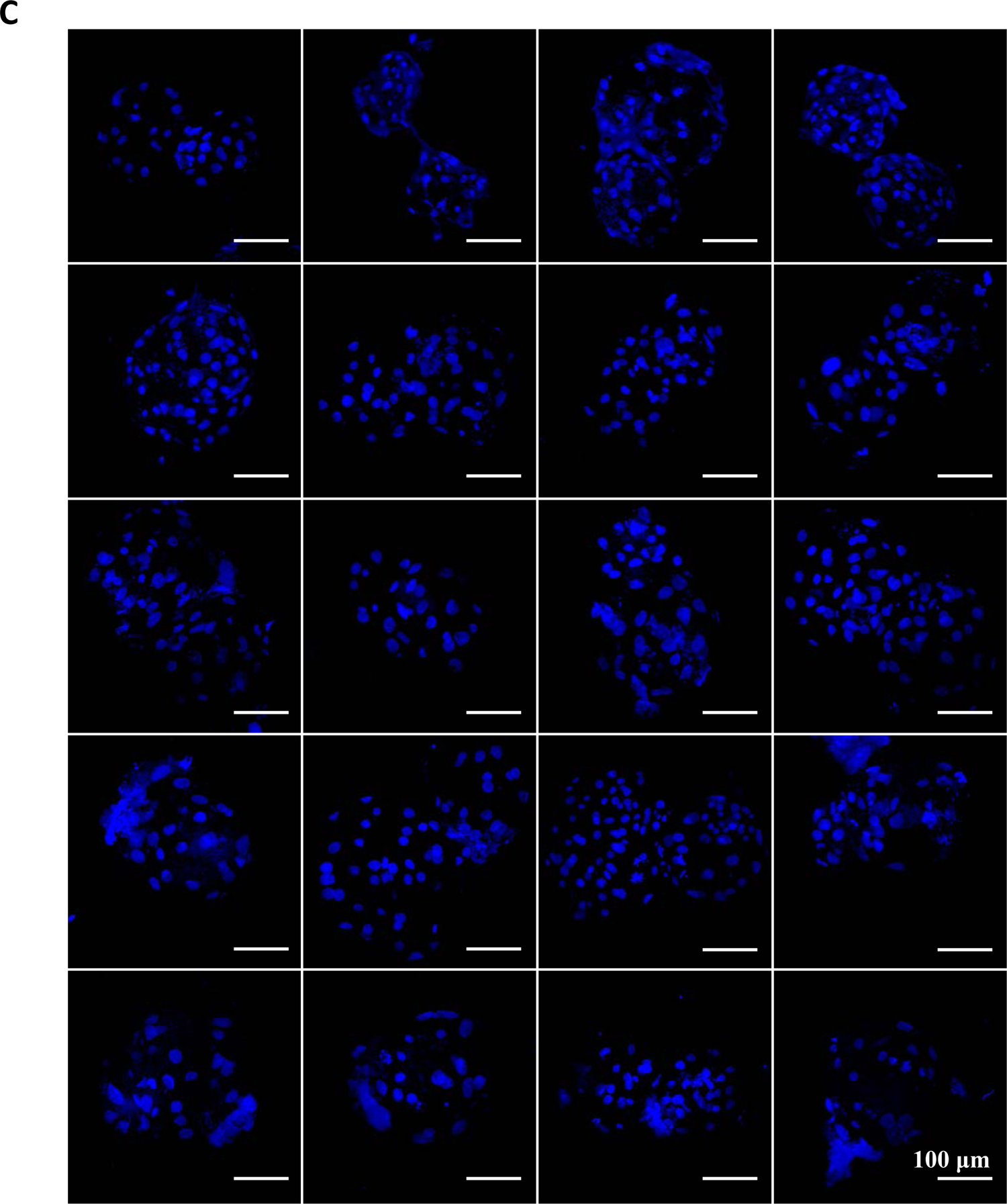
Confocal images for nucleus of untreated, Dox-treated, and Dox+4-OHT treated CIRKO-p*POU5F1* blastocyst.

**Figure 6-figure supplement 5.**
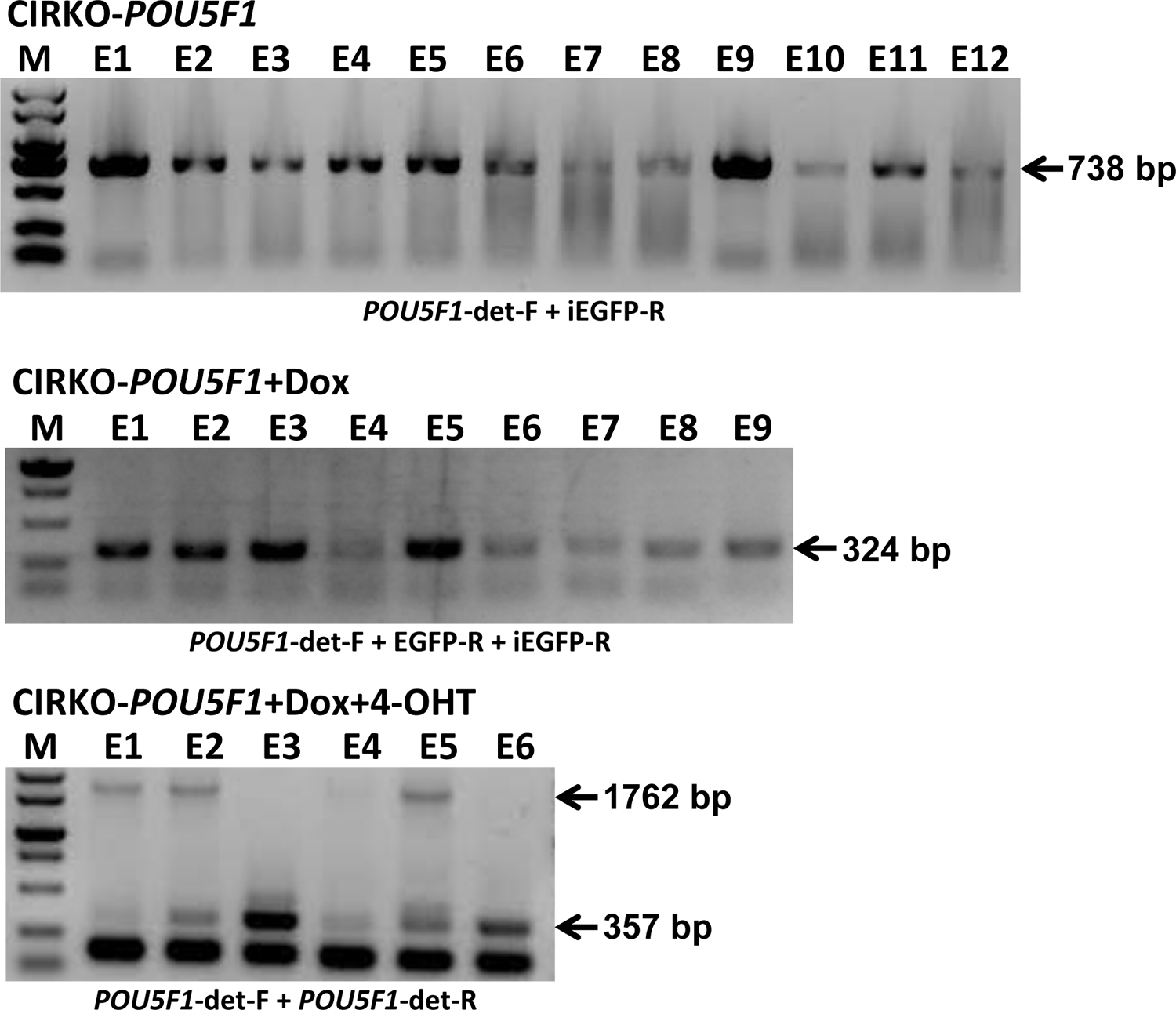
PCR analysis of p*POU5F1* locus in individual untreated, Dox-treated, and Dox+4-OHT treated CIRKO-p*POU5F1* blastocyst.

**Supplemental Table 1.**
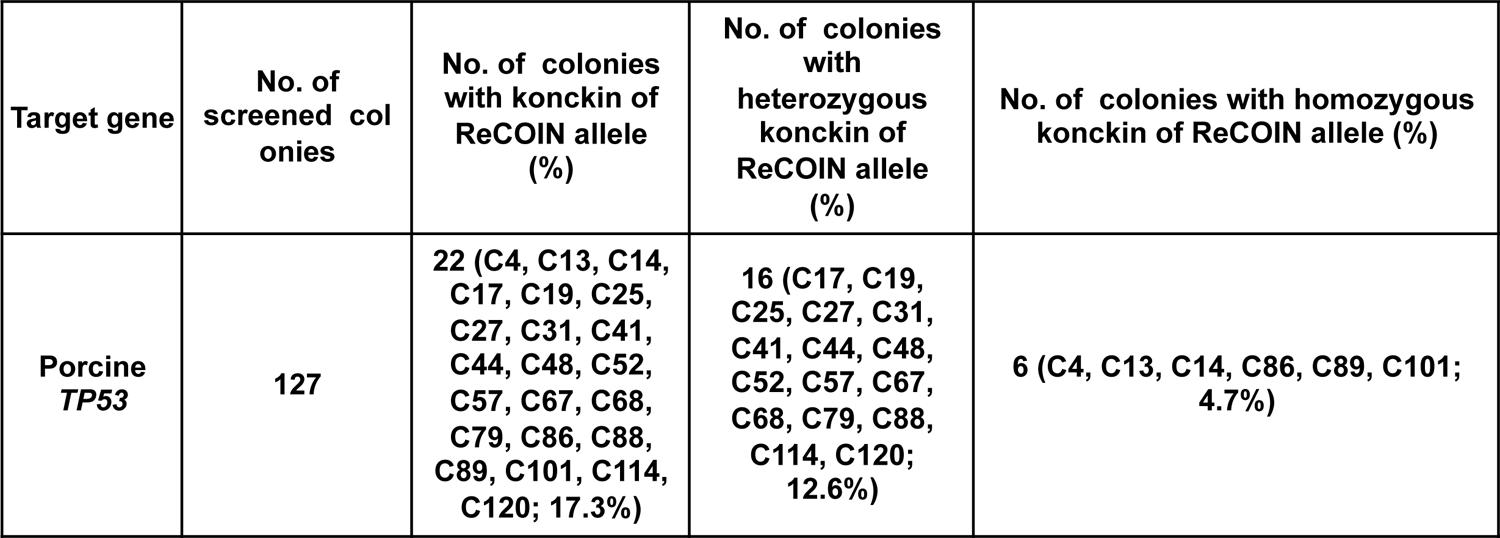
Summary of screening the ReCOIN-integrated colonies in endogenous p*TP53* gene.

**Supplemental Table 2.** Summary of generagon of cloned pigs with ReCOIN allele at endogenous porcine *TP53* gene.

| Target gene | Cell pools | Transferred embryos | No. of recipients | No. of pregnancies (%) | No. of cloned piglets | No. of piglets carrying ReCOIN allele (%) |
| --- | --- | --- | --- | --- | --- | --- |
| *Porcine TP53* | C13, C79, C86, C114 | 785 | 4 (017208, 303306, 305502, 071706) | 1 (017208; 25%) | 4 (017208-2, 017208-4, 017208-6, 017208-8) | 3 (017208-2, 017208-6, 017208-8; 75%) |

**Supplemental Table 3.** Summary of screening the CIRKO-integrated colonies in endogenous p*TP53*, p*POU5F1* and p*LMNA* gene.

| Target genes | No. of screened colonies | No. of colonies with konck in of TRE3G-controlled Cre and Flpo expression cassette at the Hipp11 locus (%) | No. of colonies with konck in of CIRKO allele at target genes (%) | No. of colonies with double knock in (%) |
| --- | --- | --- | --- | --- |
| Hipp11 and TP53 | 61 | 21 (3, 6, 8, 13, 16, 18, 21, 22, 25, 29, 31, 34, 35, 38, 39, 40, 41, 46, 54, 57, 58; 34.4%) | 12 (3, 8, 13, 16, 18, 22, 26, 31, 35, 39, 45, 60; 19.7%) | 9 (3, 8, 13, 16, 18, 22, 31, 35, 39; 14.8%) |
| Hipp11 and POU5F1 | 36 | 9 (3, 12, 13, 15, 19, 28, 32, 35, 36; 25.0%) | 23 (1, 3, 5, 7, 9, 12, 13, 15, 17, 19, 20, 21, 22, 23, 25, 26, 27, 28, 29, 32, 33, 35, 36; 63.9%) | 9 (3, 12, 13, 15, 19, 28, 32, 35, 36; 25.0%) |
| Hipp11 and LMNA | 21 | 10 (2, 4, 5, 8, 9,10, 12, 13, 15, 16; 47.6%) | 13 (2, 3, 4, 5, 6, 8, 9, 10, 12, 13, 15, 16, 19; 61.9%) | 9 (2, 4, 5, 8, 10, 12, 13, 15, 16; 42.9%) |

**Supplemental Table 4.** Blastocyst rate of reconstructed embryos from untreated, Dox-treated, and Dox+4-OHT treated CIRKO-p*POU5F1* PFFs.

| Treatment | No. of Experiments | No. of SCNT embryos | No. of Blastocysts | Blastocyst rate (%) |
| --- | --- | --- | --- | --- |
| Blank | 3 | 104 | 26 | 25.00000 |
|  |  | 95 | 14 | 14.73684 |
|  |  | 62 | 10 | 16.12903 |
| Dox | 3 | 110 | 14 | 12.72727 |
|  |  | 103 | 18 | 17.47573 |
|  |  | 91 | 20 | 21.97802 |
| Dox, 4-OHT | 3 | 115 | 19 | 16.52174 |
|  |  | 82 | 8 | 9.756097 |
|  |  | 81 | 13 | 16.04938 |

**Supplemental Table 5.**
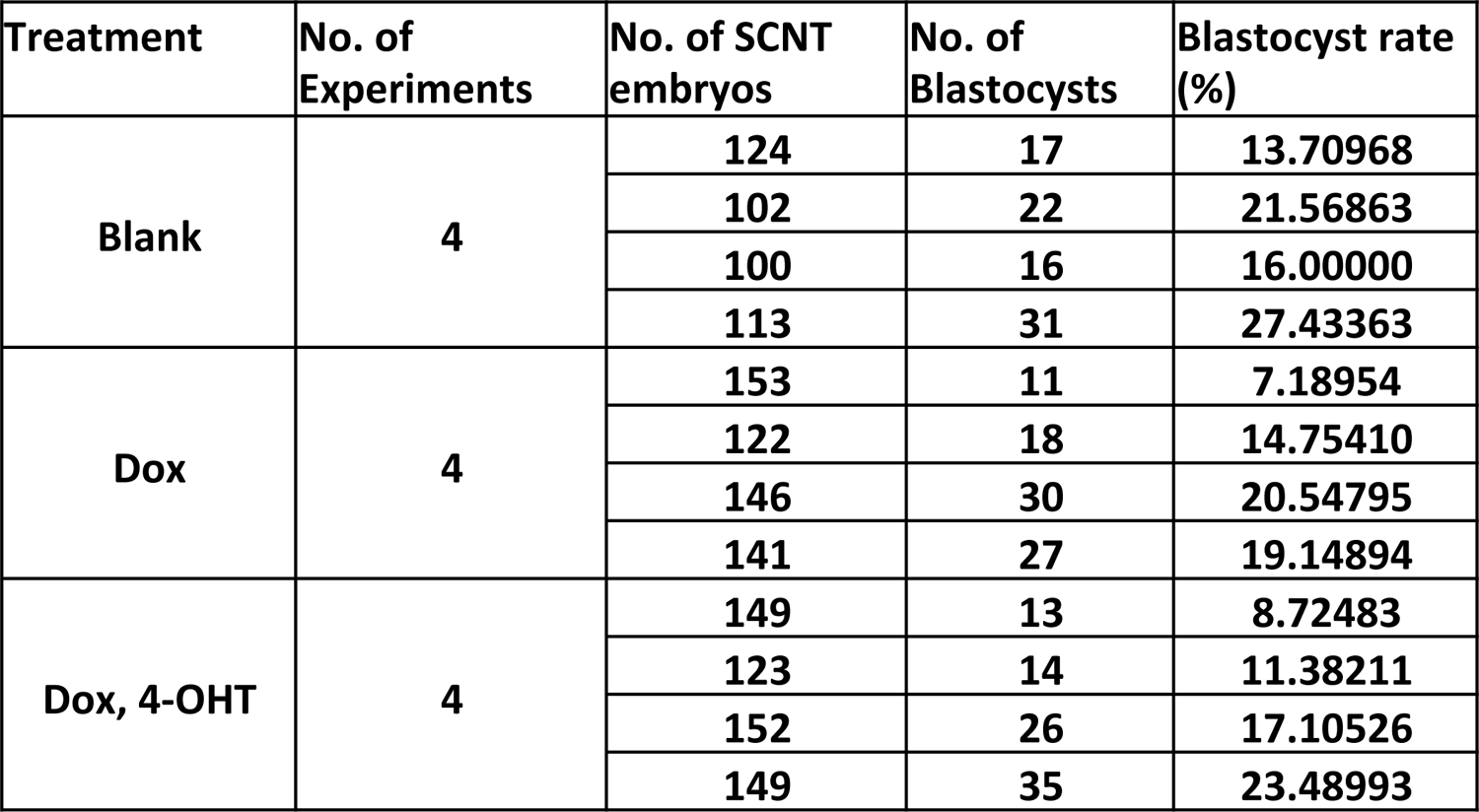
Blastocyst rate of untreated, Dox-treated, and Dox+4-OHT treated CIRKO-p*POU5F1* reconstructed embryos.

**Supplemental Table 6.**
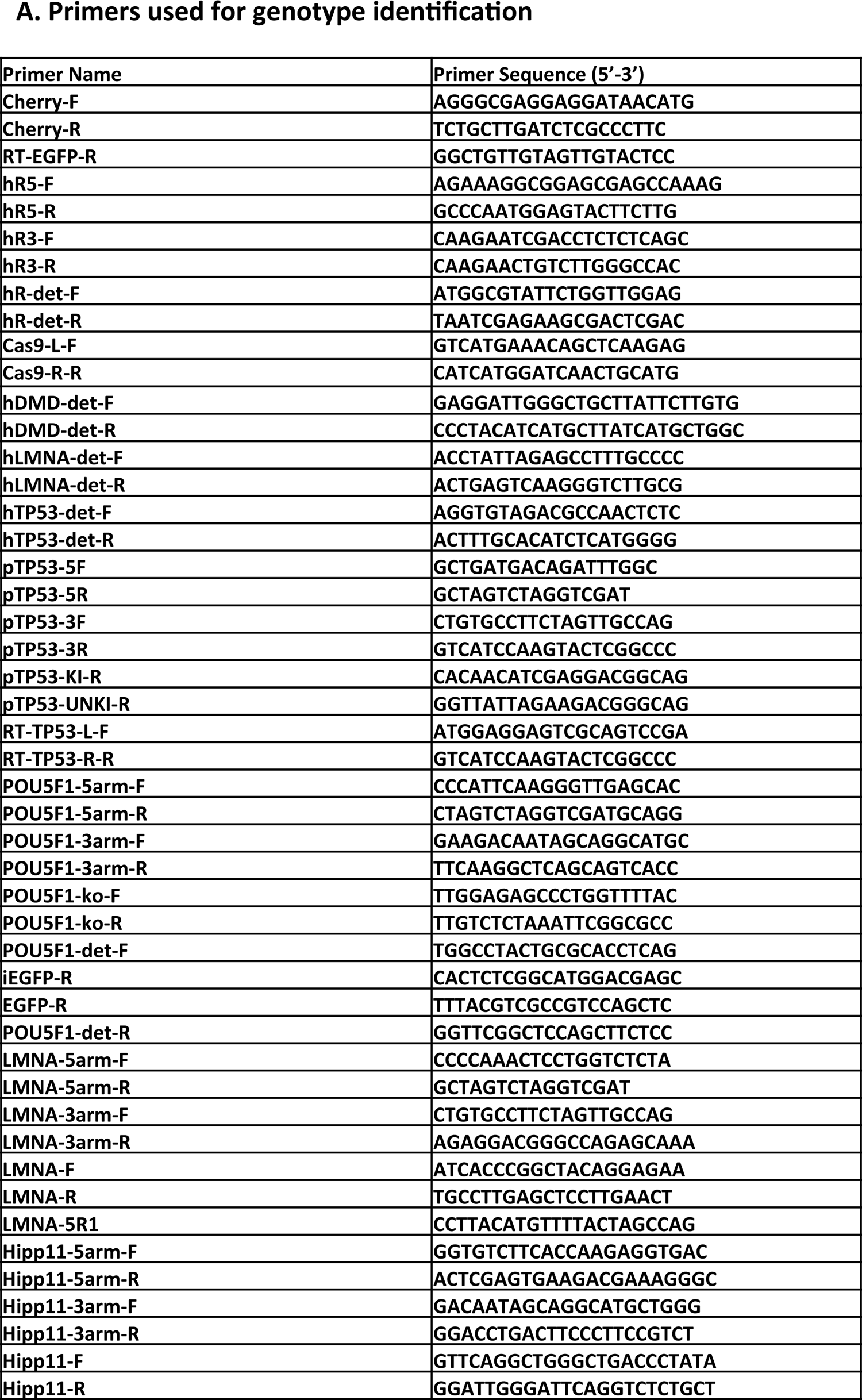
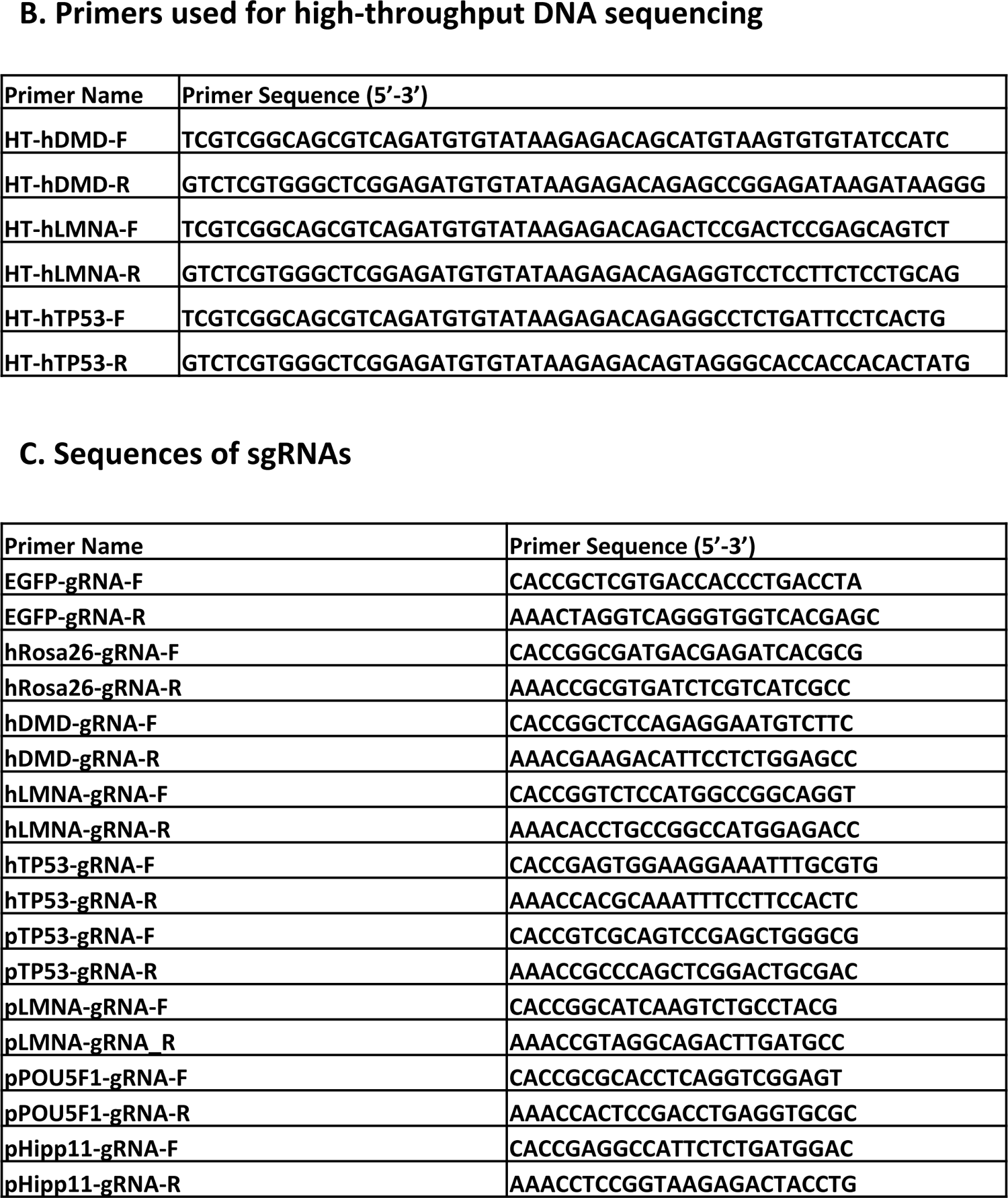
Primer sequences used in the study

## Notes

### Competing Interest Statement

The authors have declared no competing interest.

## References

1. Andersson-Rolf A, Mustata R C, Merenda A, Kim J, Perera S, Grego T, Andrews K, Tremble K, Silva J C, Fink J, Skarnes W C, Koo B K. 2017. One-step generation of conditional and reversible gene knockouts. Nat Methods 14: 287–289. doi: 10.1038/nmeth.4156.

2. Chen M, Shi H, Gou S, Wang X, Li L, Jin Q, Wu H, Zhang H, Li Y, Wang L, Li H, Lin J, Guo W, Jiang Z, Yang X, Xu A, Zhu Y, Zhang C, Lai L, Li X. 2021. In vivo genome editing in mouse restores dystrophin expression in Duchenne muscular dystrophy patient muscle fibers. Genome Med 13: 57. doi: 10.1186/s13073-021-00876-0.

3. Chew W L. 2018. Immunity to CRISPR Cas9 and Cas12a therapeutics. Wiley Interdiscip Rev Syst Biol Med 10. doi: 10.1002/wsbm.1408.

4. Clement K, Rees H, Canver M C, Gehrke J M, Farouni R, Hsu J Y, Cole M A, Liu D R, Joung J K, Bauer D E, Pinello L. 2019. CRISPResso2 provides accurate and rapid genome editing sequence analysis. Nat Biotechnol 37: 224–226. doi: 10.1038/s41587-019-0032-3.

5. Cox D B, Platt R J, Zhang F. 2015. Therapeutic genome editing: prospects and challenges. Nat Med 21: 121–131. doi: 10.1038/nm.3793.

6. Davis K M, Pattanayak V, Thompson D B, Zuris J A, Liu D R. 2015. Small molecule-triggered Cas9 protein with improved genome-editing specificity. Nat Chem Biol 11: 316–318. doi: 10.1038/nchembio.1793.

7. Donehower L A, Lozano G. 2009. 20 years studying p53 functions in genetically engineered mice. Nat Rev Cancer 9: 831–841. doi: 10.1038/nrc2731.

8. Economides A N, Frendewey D, Yang P, Dominguez M G, Dore A T, Lobov I B, Persaud T, Rojas J, McClain J, Lengyel P, Droguett G, Chernomorsky R, Stevens S, Auerbach W, Dechiara T M, Pouyemirou W, Cruz J M, Jr., Feeley K, Mellis I A, Yasenchack J, Hatsell S J, Xie L, Latres E, Huang L, Zhang Y, Pefanis E, Skokos D, Deckelbaum R A, Croll S D, Davis S, Valenzuela D M, Gale N W, Murphy A J, Yancopoulos G D. 2013. Conditionals by inversion provide a universal method for the generation of conditional alleles. Proc Natl Acad Sci U S A 110: E3179–3188. doi: 10.1073/pnas.1217812110.

9. Enache O M, Rendo V, Abdusamad M, Lam D, Davison D, Pal S, Currimjee N, Hess J, Pantel S, Nag A, Thorner A R, Doench J G, Vazquez F, Beroukhim R, Golub T R, Ben-David U. 2020. Cas9 activates the p53 pathway and selects for p53-inactivating mutations. Nat Genet 52: 662–668. doi: 10.1038/s41588-020-0623-4.

10. Fan N, Lai L. 2013. Genetically modified pig models for human diseases. J Genet Genomics 40: 67–73. doi: 10.1016/j.jgg.2012.07.014.

11. Feil R, Wagner J, Metzger D, Chambon P. 1997. Regulation of Cre recombinase activity by mutated estrogen receptor ligand-binding domains. Biochem Biophys Res Commun 237: 752–757. doi: 10.1006/bbrc.1997.7124.

12. Frum T, Halbisen M A, Wang C, Amiri H, Robson P, Ralston A. 2013. Oct4 cell-autonomously promotes primitive endoderm development in the mouse blastocyst. Dev Cell 25: 610–622. doi: 10.1016/j.devcel.2013.05.004.

13. Genschel J, Schmidt H H. 2000. Mutations in the LMNA gene encoding lamin A/C. Hum Mutat 16: 451–459. doi: 10.1002/1098-1004(200012)16:6<451::AID-HUMU1>3.0.CO;2-9.

14. Housden B E, Muhar M, Gemberling M, Gersbach C A, Stainier D Y, Seydoux G, Mohr S E, Zuber J, Perrimon N. 2017. Loss-of-function genetic tools for animal models: cross-species and cross-platform differences. Nat Rev Genet 18: 24–40. doi: 10.1038/nrg.2016.118.

15. Imperatore F, Maurizio J, Vargas Aguilar S, Busch C J, Favret J, Kowenz-Leutz E, Cathou W, Gentek R, Perrin P, Leutz A, Berruyer C, Sieweke M H. 2017. SIRT1 regulates macrophage self-renewal. EMBO J 36: 2353–2372. doi: 10.15252/embj.201695737.

16. Jin Q, Yang X, Gou S, Liu X, Zhuang Z, Liang Y, Shi H, Huang J, Wu H, Zhao Y, Ouyang Z, Zhang Q, Liu Z, Chen F, Ge W, Xie J, Li N, Lai C, Zhao X, Wang J, Lian M, Li L, Quan L, Ye Y, Lai L, Wang K. 2022. Double knock-in pig models with elements of binary Tet-On and phiC31 integrase systems for controllable and switchable gene expression. Sci China Life Sci. doi: 10.1007/s11427-021-2088-1.

17. Lai L, Prather R S. 2003. Production of cloned pigs by using somatic cells as donors. Cloning Stem Cells 5: 233–241. doi: 10.1089/153623003772032754.

18. Leibowitz M L, Papathanasiou S, Doerfler P A, Blaine L J, Sun L, Yao Y, Zhang C Z, Weiss M J, Pellman D. 2021. Chromothripsis as an on-target consequence of CRISPR-Cas9 genome editing. Nat Genet 53: 895–905. doi: 10.1038/s41588-021-00838-7.

19. Lewandoski M. 2001. Conditional control of gene expression in the mouse. Nat Rev Genet 2: 743–755. doi: 10.1038/35093537.

20. Liu K I, Ramli M N, Woo C W, Wang Y, Zhao T, Zhang X, Yim G R, Chong B Y, Gowher A, Chua M Z, Jung J, Lee J H, Tan M H. 2016. A chemical-inducible CRISPR-Cas9 system for rapid control of genome editing. Nat Chem Biol 12: 980–987. doi: 10.1038/nchembio.2179.

21. Lu H, Liu J, Feng T, Guo Z, Yin Y, Gao F, Cao G, Du X, Wu S. 2021. A HIT-trapping strategy for rapid generation of reversible and conditional alleles using a universal donor. Genome Res 31: 900–909. doi: 10.1101/gr.271312.120.

22. Lundin A, Porritt M J, Jaiswal H, Seeliger F, Johansson C, Bidar A W, Badertscher L, Wimberger S, Davies E J, Hardaker E, Martins C P, James E, Admyre T, Taheri-Ghahfarokhi A, Bradley J, Schantz A, Alaeimahabadi B, Clausen M, Xu X, Mayr L M, Nitsch R, Bohlooly Y M, Barry S T, Maresca M. 2020. Development of an ObLiGaRe Doxycycline Inducible Cas9 system for pre-clinical cancer drug discovery. Nat Commun 11: 4903. doi: 10.1038/s41467-020-18548-9.

23. Martins C P, Brown-Swigart L, Evan G I. 2006. Modeling the therapeutic efficacy of p53 restoration in tumors. Cell 127: 1323–1334. doi: 10.1016/j.cell.2006.12.007.

24. Maynard L H, Humbert O, Peterson C W, Kiem H P. 2021. Genome editing in large animal models. Mol Ther. doi: 10.1016/j.ymthe.2021.09.026.

25. Meurens F, Summerfield A, Nauwynck H, Saif L, Gerdts V. 2012. The pig: a model for human infectious diseases. Trends Microbiol 20: 50–57. doi: 10.1016/j.tim.2011.11.002.

26. Neff E P. 2019. Cancer modeling thinks big with the pig. Lab Anim (NY) 48: 75–78. doi: 10.1038/s41684-019-0246-5.

27. Nichols J, Zevnik B, Anastassiadis K, Niwa H, Klewe-Nebenius D, Chambers I, Scholer H, Smith A. 1998. Formation of pluripotent stem cells in the mammalian embryo depends on the POU transcription factor Oct4. Cell 95: 379–391. doi: 10.1016/s0092-8674(00)81769-9.

28. Park J, Bae S, Kim J S. 2015. Cas-Designer: a web-based tool for choice of CRISPR-Cas9 target sites. Bioinformatics 31: 4014–4016. doi: 10.1093/bioinformatics/btv537.

29. Robles-Oteiza C, Taylor S, Yates T, Cicchini M, Lauderback B, Cashman C R, Burds A A, Winslow M M, Jacks T, Feldser D M. 2015. Recombinase-based conditional and reversible gene regulation via XTR alleles. Nat Commun 6: 8783. doi: 10.1038/ncomms9783.

30. Schwenk F, Kuhn R, Angrand P O, Rajewsky K, Stewart A F. 1998. Temporally and spatially regulated somatic mutagenesis in mice. Nucleic Acids Res 26: 1427–1432. doi: 10.1093/nar/26.6.1427.

31. Utomo A R, Nikitin A Y, Lee W H. 1999. Temporal, spatial, and cell type-specific control of Cre-mediated DNA recombination in transgenic mice. Nat Biotechnol 17: 1091–1096. doi: 10.1038/15073.

32. Vogelstein B, Lane D, Levine A J. 2000. Surfing the p53 network. Nature 408: 307–310. doi: 10.1038/35042675.

33. Vousden K H, Lu X. 2002. Live or let die: the cell’s response to p53. Nat Rev Cancer 2: 594–604. doi: 10.1038/nrc864.

34. Wu H, Liu Q, Shi H, Xie J, Zhang Q, Ouyang Z, Li N, Yang Y, Liu Z, Zhao Y, Lai C, Ruan D, Peng J, Ge W, Chen F, Fan N, Jin Q, Liang Y, Lan T, Yang X, Wang X, Lei Z, Doevendans P A, Sluijter J P G, Wang K, Li X, Lai L. 2018. Engineering CRISPR/Cpf1 with tRNA promotes genome editing capability in mammalian systems. Cell Mol Life Sci 75: 3593-3607. doi: 10.1007/s00018-018-2810-3.

35. Xu S, Kim J, Tang Q, Chen Q, Liu J, Xu Y, Fu X. 2020. CAS9 is a genome mutator by directly disrupting DNA-PK dependent DNA repair pathway. Protein Cell 11: 352–365. doi: 10.1007/s13238-020-00699-6.

36. Yarmolinsky M, Hoess R. 2015. The Legacy of Nat Sternberg: The Genesis of Cre-lox Technology. Annu Rev Virol 2: 25–40. doi: 10.1146/annurev-virology-100114-054930.

37. Zhang J, Khazalwa E M, Abkallo H M, Zhou Y, Nie X, Ruan J, Zhao C, Wang J, Xu J, Li X, Zhao S, Zuo E, Steinaa L, Xie S. 2021. The advancements, challenges, and future implications of the CRISPR/Cas9 system in swine research. J Genet Genomics 48: 347–360. doi: 10.1016/j.jgg.2021.03.015.

